# Phenotypic and Genotypic Characterization of the Population of *Phytophthora infestans* in Brazil

**DOI:** 10.1101/2025.09.03.673969

**Authors:** Mariana G. Silva, Gabriel Alves, Letícia C. Araújo, Saulo A. S. Oliveira, Eduardo S. G. Mizubuti

**Affiliations:** Departamento de Fitopatologia, Universidade Federal de Viçosa, 26570-900 Viçosa, MG, Brazil; Embrapa Mandioca e Fruticultura, Cruz das Almas, BA, Brazil

## Abstract

Recently, phenotypic variations and differences in virulence have been observed among Brazilian isolates of *P*. *infestans*. Analysis of phenotypic variability can be useful to understand the pathogen population and how variants may affect late blight epidemics. Thus, this study aimed to characterize the population of *P*. *infestans* in Brazil using mating type, mitochondrial DNA haplotype, microsatellite genotype, and mefenoxam sensitivity. Isolates of the A2 mating type from tomato are reported for the first time. Isolates from potato plants were either of A1 or A2 mating types. The Ia and Ib mitochondrial DNA haplotypes were detected in Brazil. The Ia is associated with isolates from potato and the Ib with isolates obtained from potato and tomato. All tomato isolates were of a new genotype, SA_1_A2, while the EU_37_A2 and EU_2_A1 genotypes were detected among isolates sampled from potato fields. The sensitivity to mefenoxam varied: sensitive, intermediately insensitive, and insensitive isolates are present in potato and tomato crops. The population of *P*. *infestans* in Brazil remains structured according to host, but genotypes associated with potato or tomato plants seem to vary periodically.

## Introduction

Populations of *Phytophthora infestans* (Mont.) de Bary have been studied and characterized for many decades, in several parts of the globe, trying to uncover epidemiologically-relevant evolutionary processes at different time scales <u>(Huang et al. 2024; Coomber et al. 2025)</u>. The importance of this pathogen explains the large number of research groups worldwide dedicated to studying population genetics, ecology, and evolution, with the final goal of improving knowledge about late blight epidemiology, management strategies and, ultimately, reducing yield losses.

In Brazil, the first study of the population of *P. infestans* using molecular markers was performed in the late 1980’s. Back then, isolates of A1 mating type were reported to be associated with tomato, while A2 isolates were associated with potato (Brommonschenkel 1988). Based on isolates obtained in the late 1980’s, two clonal lineages of *P. infestans* were confirmed to be present in Brazil: US-1 (A1 mating type) and BR-1 (A2), associated with tomato and potato plants, respectively (Goodwin et al. 1994). Despite the presence of individuals of the two mating types for at least a decade in the fields, sexual recombination was not detected in the Southern, Southeast, and Central-West of Brazil (Reis et al. 2003, 2006). The association of each of the mating types with different hosts may have prevented sexual reproduction and also suggested host preference according to this marker. Later studies demonstrated that the population of *P. infestans* remained structured by host (Reis et al. 2003, 2006) and host specialization was reported (Reis et al. 2003). Although isolates from one host can infect the other, the host-genotype association resulted in differences in pathogen virulence: the US-1 isolates were more virulent to tomato while BR-1 isolates were more virulent to potato (Suassuna et al. 2004).

Subsequently, new studies revealed important changes in the pathogen population in Brazil. For the first time, individuals of the US-1 genotype and A1 mating type were found causing late blight on potato in the Southern region (Santana et al. 2013). In Rio Grande do Sul state, individuals of both mating types were found co-occurring in potato fields, prompting for the possibility of sexual recombination (Santana et al. 2013). Additionally, self-fertile isolates of *P*. *infestans* were reported for the first time in Brazil, i.e. isolates able to form oospores by themselves and when paired with isolates of either mating type <u>(Smart et al. 2000)</u>. Even with the presence of individuals of A1 and A2 mating types now occurring simultaneously on potato fields, evidence of sexual reproduction happening in Brazil was only found in the Southern region between 2003 and 2005 (Oliveira 2010; Santana et al. 2013).

In the latest studies conducted in Brazil, self-fertile isolates were reported in fields sampled in Paraná state in 2011 and 2012 (Casa-Coila et al. 2017). Also, oospores were found in naturally infected potato leaves in four fields sampled in São Paulo, Minas Gerais, and Paraná states (Zanotta 2019). Additionally, EU_2_A1 was reported as the most frequently found genotype in potato fields in these states in the 2015 season (Zanotta 2019). This suggests that EU_2_A1 could have displaced BR-1 as the predominant lineage associated with potato in Brazil (Zanotta 2019).

The dynamics of the *P*. *infestans* population over the past decades in Brazil indicates important changes of evolutionary and epidemiological consequences. Migration of new lineages or even rare events of sexual recombination may have altered the structure of the population and interfered in the efficiency of management strategies. Therefore, populations of *P. infestans* should be monitored on a regular basis to detect alterations that may result in severe epidemics.

Phenotypic variations and differences in virulence among isolates of *P*. *infestans* have been observed in Brazil, leading us to hypothesize that there was a shift in the population of *P*. *infestans* in Brazil. Thus, this study aimed to (i) characterize the current Brazilian population of *P*. *infestans* in mating type, mitochondrial DNA haplotype, 12-plex SSR genotype, and mefenoxam sensitivity, and (ii) to conduct a phylogenetic analysis of the isolates sampled from tomato and potato fields to check for any evidence of host association.

## Material and Methods

### Collection of *P*. *infestans* isolates

Leaves, fruits (tomato) and stems of potato and tomato plants with typical late blight lesions were collected from fields located in Bahia (BA), Minas Gerais (MG), Espírito Santo (ES), São Paulo (SP), Paraná (PR), Santa Catarina (SC), and Rio Grande do Sul (RS) states, from 2020 to 2024. The infected plant material was kept in paper bags until they arrived at the laboratory. The samples were incubated in a humid chamber at 18°C in the dark to induce sporangia formation. The tomato samples were separated into two groups: 1) leaves with late blight lesions only, and 2) leaves with late blight lesions and necrotic spots caused by other pathogens. The samples of the first group were used for direct isolation and leaves from the second group were used for indirect isolation procedures. The pathogen isolation by direct method was attempted by removing sporangia by means of insulin needle from sporangiophores formed on the interface between the necrotic region of a single lesion and the green tissues of plant material. Sporangia were transferred to pea-agar medium (PAM) (150 g pea, 15 g agar, 0.05 g beta-sitosterol per liter) or V8 10 % unclarified amended with rifampicin (0.02 g/L), ampicillin (0.10 g/L), and tebuconazole fungicide (Folicur®) (10 mg a.i./L) or thiabendazole fungicide (Tecto®) (10 mg a.i./L). The Petri dishes containing sporangia were kept at 18 °C in darkness for 5 to 10 days and were visually inspected at every two days under a stereomicroscope.

Colonies with coenocytic hyphae, forming limoniform and papillated sporangia on branched sporangiophores with the distinctive morphological aspects of *P*. *infestans* were selected and mycelial discs of colonies were transferred to PAM or to PARP medium (500 mL of Pea Agar, 0.125 g ampicillin, 0.05 g PCNB, and 0.005 g rifampicin) and kept at 18 °C in darkness for 14 days. To obtain monozoosporic cultures, pure cultures were washed with 8 ml of cold sterile deionized water (4 °C) and the resulting sporangia suspensions were kept at 4 °C for 2 to 3 h to stimulate zoospores release. An aliquot was collected with a platinum loop, spread on the surface of PAM in a zigzag pattern, previously marked on the bottom of the Petri dish, or an aliquot of 100 μL of the zoospore suspension was spread on the surface of the water-agar medium (20 g agar per liter) by Drigalski handle. The Petri dishes were kept at 10 °C in darkness for 12 h to allow for germination of the zoospores. A germinated zoospore was selected with the help of a microscope (Olympus CX31) at 400X, transferred to PAM, and kept at 18 °C in darkness for 14 days.

For the indirect isolation method on tomato samples, small pieces of leaves (∼ 5.0 x 5.0 mm) were taken from the intersection area between the necrotic region of a single lesion and the green tissues using a sterilized scalpel. The tissue fragments were washed in 1 % sodium hypochlorite for 30 s, followed by two washes in autoclaved distilled water for 30 s each and immediately placed on filter paper to dry the excess water and then transferred to Petri dishes containing PARP medium with the abaxial part facing up. The plates were kept in an incubator at 18 °C. Colonies were visually inspected daily with the aid of a stereomicroscope (Olympus SZ61 model SZ2-ILST) at 10X.

In addition to isolation to culture medium, *P*. *infestans* isolates from tomato plants were kept as living colonies in asymptomatic tomato leaves. One leaflet of tomato samples, from each sampled area, with a single sporulating lesion of *P*. *infestans* was selected and placed in non-chlorinated water at 4 °C for 2 h to induce zoospore release. Subsequently, asymptomatic tomato leaves from greenhouse-grown plants were inoculated with 15 µL-drops of the zoospore suspension. Four to five drops were deposited per leaflet. The inoculated leaves were kept in a humid chamber in an incubator at 18 °C in the dark. Lesions formed in inoculated leaflets were subsequently and periodically transferred to asymptomatic leaves at every 10 days.

#### DNA extraction

Mycelial discs of the isolates were transferred to pea broth (120 g pea/L) and kept at 18 °C in darkness, for 8 to 15 days (Lindqvist-Kreuze et al. 2020). After colony development, mycelia were washed twice with sterile distilled water, completely dried at a laminar flow cabinet, and kept at -20 °C until DNA extraction.

The DNA was extracted with the Wizard® Genomic DNA Purification kit (Promega Corporation). The quality and quantity of DNA were accessed using Nanodrop, visualized in 1 % agarose electrophoresis gel, and its concentration was standardized at 20 ng/µL for genotypic characterization and 100 ng/µL for characterization of mating type, mitochondrial DNA, and for sequencing of PCR-amplicons of partial genomic regions.

#### Mating type characterization

The mating type of each isolate was characterized using a PCR-based method and the results were confronted with a subset of pairings made on culture medium. The mating type classification as A1 or A2 was done by PCR using standard DNA sent by Dr. David Cooke (James Hutton Institute - JHI). Subsequently, random isolates were paired on culture medium (PAM) with known testers (A1 = Pi94; and A2 = Pi88).

For the PCR-based identification of mating type primers W16-1 (AACACGCACAAGGCATATAAATGTA) and W16-2 (GCGTAATGTAGCGTAA-CAGCTCTC) were used (Judelson et al. 1995). The PCR was conducted in a 15 µL reaction mixture with 2.5 µL 5X Green GoTaq® Reaction Buffer (Promega Corporation), 0.25 µL MgCl_2_, 0.25 µL, dNTPs, 0.1 µL GoTaq® Polymerase, 0.2 µL each primer, 9.5 µL of sterile Milli-Q water, and 2.0 µL DNA. The PCR conditions were as described by Brylińska et al. (2018). After PCR amplification, amplicons were subjected to enzymatic digestion with restriction endonuclease from *Bacillus sphaericus* (BshFI) (Jena Bioscience) that recognizes and cuts the 5’-GGCC-3′ sequence (Vlatakis et al. 1989) on mixture contained 10 µL of PCR product, 1 µL of the enzyme, 5 µL of universal buffer, and 34 µL of sterile Milli-Q water. The digestion reaction conditions recommended by the manufacturer were: incubation at 37 °C for 10 min and inactivation at 80 °C for 20 min. The PCR and enzymatic digestion products were separated and visualized in 1 % and 2 % agarose electrophoresis gel, respectively. The presence of fragments of approximately 100, 500, and 600 bp identify the A1 mating type and the presence of amplicons of near to 100 and 500 bp identify the A2 mating type.

The validation of the mating-type primers was conducted with randomly selected isolates that were paired with A1 and A2 tester isolates previously identified by the PCR-based method. A strip (∼ 3 cm length x 0.5 cm width) of the culture medium and mycelia of the “query” (unknown) isolate from 7 to 15-day-old colonies was placed in the middle of a Petri dish containing PAM. A mycelial strip of the A1 tester isolate was placed approximately 3 cm away and to the right of the centered-placed unknown isolate and another strip of the A2 tester was placed 3 cm away and to the left of the strip of the query isolate. Petri dishes were kept at 18 °C in darkness for 3 to 4 weeks and the presence of oospores was visualized in a microscope (Olympus CX31) at 400X. If the query isolate formed oospores with the A1 tester it was identified as A2 mating type, if the isolate formed oospores with the A2 tester it was identified as A1 mating type, if the isolate produced oospores with both testers it was deemed as A1/A2, and if the isolate produced oospores by itself it was identified as self-fertile (SF). The PCR-based identification of mating type and pairings on culture medium were conducted twice for all isolates.

#### Mitochondrial DNA haplotypes identification

The mitochondrial DNA haplotypes were identified using a set of primers that anchor in the P2 and P4 gene regions (Griffith and Shaw 1998). The PCR was conducted in a 15 µL reaction mixture with 1.25 µL 10X Reaction Buffer Complete (Cellco Biotech), 1.0 µL MgCl_2_, 0.25 µL, dNTPs, 0.1 µL Taq DNA Polymerase, 0.5 µL each primer Fwd/Rev, 9.0 µL of sterile Milli-Q water, and 2.0 µL DNA. The PCR reactions were performed under the following conditions: one cycle of 94 °C for 90 s, followed by 40 cycles of 94 °C for 40 s, 55 °C for 60 s, and 72 °C for 90 s, plus a final extension period of 72 °C for 5 min.

The expected sizes of the PCR products were 1.2 kb for P2 and 0.96 kb for P4. The amplification products of the P2 region were digested with *MspI* restriction enzyme (New England Biolabs) that recognizes and cuts the 5’-CCGG-3’ site and the amplification products of P4 region were digested with *EcoRI* restriction enzyme (Cellco Biotech) that recognizes and cuts the 5’–GAATTC–3’ sequence, according to manufacturer protocols. Fifteen µL of the digested product was stained with GelRed^TM^ (Biotium, Inc.), separated by electrophoresis on 2 % agarose gels, visualized using standard techniques, and the haplotypes were determined as previously described by Griffith & Shaw (1998).

#### Mefenoxam sensitivity

Technical grade metalaxyl-M or mefenoxam (1000.0 μg/mL) dispersible oil suspension was provided by Syngenta. An aliquot of the technical grade was used as the stock solution and stored at 4 °C in darkness. Serial dilutions were prepared from the stock solution (1000.0 μg/mL) to reach the final concentrations of 100.0, 50.0, 10.0, and 5.0 μg/mL of mefenoxam.

Mefenoxam sensitivity was assessed for isolates obtained from potato using the agar method, with slight modifications (Reis et al. 2005). The isolates were grown in PAM for 15 days and transferred to four plates (four replicates) with pea glucose agar (150 g pea, 5 g glucose, and 15 g agar per liter) amended with mefenoxam (5.0 or 100.0 μg/mL) or DMSO 0.1 % (v/v) (control). After 9 days, the mycelial growth was assessed by measuring colony diameter in two perpendicular directions using a digital caliper. The test was done twice for each isolate.

Mefenoxam sensitivity of tomato isolates was assessed using the leaf disc method (Fungicide Resistance Action Committee (FRAC) 2017). The abaxial side of tomato leaves was sprayed with 3 mL of mefenoxam suspension (100.0 and 5.0 μg/mL of i.a.) using a Devilbiss sprayer connected to an air compressor at 15 PSI (Air Plus Schulz MS2.3). After the mefenoxam application, leaves were dried in a laminar flow for 20 min, cut with a cork borer, and immediately placed in wells of 24-well plastic plates containing 1.5 mL of 0.4 % water-agar. After 24 h of mefenoxam spray, a 30 μL-drop sporangia suspension (2 x 10^4^ sporangia/mL), kept for 2 h at 4°C to induce zoospore release, was placed in a center of leaf discs. The suspension was prepared by washing sporangia of *P*. *infestans* isolates cultivated on detached leaves of tomato cv. Santa Clara with sterilized non-chlorinated water. The 24-well plastic plates were kept at 18 °C under 18 h of light alternated with 6 h of darkness for 10 to 15 days. Two days after inoculation, the remaining drop of sporangia suspension was dried with tissue paper. At the end of the incubation period, sporulation was visually observed using a stereomicroscope (Olympus SZ61 model SZ2-ILST) at 10X, and grades were assigned according to the scale described by Sozzi (1991). Each treatment was assessed using four leaf discs. The experiment was conducted twice for each isolate.

#### SSR genotyping

The isolates were genotyped by one-step SSR multiplex (Li et al. 2013) using the 12 loci. DNA of eight standard genotypes was kindly provided by Dr. David Cooke (JHI) in Whatman FTA cards. The FTA cards were prepared to be used in the PCR multiplex by (a) cutting the cards with mini punching (2.0 mm); (b) adding 150 μL of QIAcard FTA Wash Buffer (Qiagen), shaking slowly in the vortex for 3 min, and removing the reagent with a pipette, (c) repeating step “b”; (d) adding 150 μL of TE-1 Buffer (10 mM Tris-HCL, 0.1 mM EDTA, pH 8.0), shaking slowly in the vortex for 3 min, and removing the reagent with a pipette; (e) repeating step “d”; and (f) air drying disc for 2 h at room temperature.

The PCR was conducted in a 13.5 μL total volume and the protocol was adapted from Euroblight (Cooke et al. 2010). The protocol consisted of 4.81 μL of water, 6.25 μL of 2X Type-IT Multiplex PCR Mix (Qiagen), 1.0 μL of Primer Mix 1 (Cooke et al. 2010), 0.3 μL Pi4B forward and reverse (Fwd/Rev) primer (10 µM), 0.4 μL SSR8 Fwd/Rev primer (10 µM), 0.2 μL D13 Fwd/Rev primer (10 µM), 0.08 μL Pi04 Fwd/Rev primer (4.7 µM), and 2 µL of DNA. The PCR conditions were the same as described by Li et al. (2013).

The fragment analysis was done at Macrogen Inc. The allele patterns were manually sized with the GeneMaker v3.0.1 (SoftGenetics). In the first run, the standard genotypes were analyzed, and the sizes of alleles were adjusted and calibrated according to Li et al. (2013) (Table S1).

#### Temporal analysis

The current Brazilian population was compared to the isolates collected in Brazil from 1998 to 2010 (“old population”) (Oliveira 2010). The data of allele patterns of the isolates from 1998 to 2010 were obtained (Oliveira 2010) and the analysis was done using the data of six SSR loci common to all isolates: Pi02, Pi04, Pi63, D13, Pi70, and G11 (Knapova and Gisi 2002; Lees et al. 2006).

#### Phylogenetic study

The phylogenetic relation between *P*. *infestans* isolated from tomato and potato was investigated by sequencing of Beta-tubulin (*Btub*), TigA gene fusion protein (*TigA*), Internal transcribed spacer (*ITS*), and Mitochondrial cytochrome C oxidase subunit I (*CoxI*) loci (Coomber et al. 2023).

The primer set used to amplify the *Btub* locus was Btub_F1 and Btub_R1 (Blair et al. 2008; Kroon et al. 2004) and the PCR conditions were one cycle at 95°C for 3 min, followed by 40 cycles of 95 °C for 30 s, 64.9 °C for 30 s, and 72 °C for 1 min, plus a final extension period of 72 °C for 15 min. The *TigA* locus was amplified by two primer sets: Tig_for and Tig_rev, and G3PDH_for and G3PDH_rev (Blair et al. 2008). To amplify the primer set Tig_for/Tig_rev the PCR conditions were: one cycle of 95 °C for 3 min, followed by 40 cycles of 95 °C for 30 s, 63.2 °C for 30 s, and 72 °C for 1 min, plus a final extension period of 72°C for 15 min; and the amplification of G3PDH_for/G3PDH_rev set was under one cycle of 95 °C for 2 min, followed by 35 cycles of 95 °C for 30 s, 64 °C for 30 s, and 72 °C for 2 min, plus a final extension period of 72 °C for 5 min. The ITS4 and ITS6 (Cooke et al. 2000) were used to amplify the *ITS* locus under PCR conditions of one cycle of 95 °C for 1.25 min, followed by 34 cycles of 95 °C for 35 s, 62 °C for 35 s, and 72 °C for 55s, and a final extension period of 72 °C for 10 min. The COXF4N and COXR4N were used (Kroon et al. 2004) to amplify the *CoxI* locus according to the conditions described by Coomber et al. (2023). The PCR products were separated and visualized in 1 % agarose electrophoresis gel, purified using ExoSAP-IT™ (Affymetrix™), and sent to Macrogen (South Korea) to be sequenced.

#### Data analysis

To classify the isolate according to their sensitivity to mefenoxam, the mycelial growth and sporulation on leaf disc of each isolate grown in the control was considered as 100 %, and the mycelial growth and sporulation on leaf disc grades with 5.0 and 100.0 μg of mefenoxam/mL were transformed in percentage in relation to the control. The isolates were classified as sensitive (growth or sporulation less than 40 % of the control at 5.0 and 100.0 μg/mL), intermediate (growth or sporulation greater than 40 % of the control at 5.0 μg/mL and less than 40 % of the control at 100.0 μg/mL), or resistant (growth or sporulation greater than 40 % of the control at both 5.0 and 100.0 μg/mL) (Therrien et al. 1993).

For genotype analyses, two datasets were constructed: 1. Brazilian isolates; and 2. Expanded dataset containing Brazilian isolates and 173 reference isolates of genotypes described in North and South America and Europe. The data for the expanded analysis were from: the October 2024 version of the dataset of Saville and Ristaino (2019) study using methods in line with the Euroblight scoring system (Euroblight UPD), the dataset of Coomber et al. (2023) study and available at *P*. *infestans* classifier (T-BAS) (Coomber et al. 2023), and the data of standard genotypes provided by Dr. David Cooke (JHI). The Bruvo’s genetic distance was calculated for the new dataset with the bruvo.dist() function and a cut-off value was estimated with function cutoff_predictor() of R (R Core Team 2024) in package *poppr* version 2.9.6 (Kamvar et al. 2014). The new dataset was updated with the data of the current population. The Bruvo’s genetic distance was calculated for the new dataset updated as described before, and the cut-off value was used to identify the genotypes of each isolate of the current population. When the Bruvo’s genetic distance was lower than the estimated cut-off value, a genotype match was considered.

To complete the identification of the genotypes, a Neighbor-Joining tree (NJ) with 1000 bootstraps and a minimum spanning network (MSN) were generated from Bruvo’s genetic distance, respectively, by functions bruvo.boot() and bruvo.msn() of the *poppr* package.

General population genetics statistics were estimated for the Brazilian population using the package *poppr*. The genotypic diversity was obtained using the poppr() function, the statistics estimated were: number of individuals observed (N), number of multilocus genotypes (MLG) observed, number of expected MLG at the smallest sample size ≥ 10 based on rarefaction (eMLG), Standard error based on eMLG (SE), Shannon-Wiener Index of MLG diversity (Shannon 1948) (H), Evenness E5 (Grünwald et al. 2003; Ludwig and Reynolds 1988; Pielou 1975) (E.5), Nei’s unbiased gene diversity (Nei 1978) (Hexp), index of association IA (Brown et al. 1980; Smith et al. 1993) (Ia), and standardized index of association 𝑟D (rbarD). An MLG histogram was generated to examine the diversity of MLGs by host, state of origin, mating type, mitochondrial DNA haplotype, and genotype by mlg.table() function. Clone correction was done individually by host, state of origin, mating type, mitochondrial haplotype, and genotype using the function clonecorrect(). Clone corrected datasets, i.e. containing one representative of each haplotype, were used to assess population structure.

Deviations from Hardy-Weinberg equilibrium (HWE) were checked by hw.test() function of *pegas* package (Paradis 2010). The population was checked for linkage disequilibrium (LD) using the function ia() with 999 permutations in package *poppr*.

The population structure was assessed using model-based Bayesian clustering in STRUCTURE v.2.3.4 (Pritchard et al. 2000). The parameters used were burn-in of 20,000 repeats, 1,000,000 Markov chain Monte Carlo (MCMC) repeats, and the admixture model was chosen. Independent runs of the model used K values from 1 to 10 with 20 iterations (runs) of each K value. The best K was estimated by the Evanno method (Evanno et al. 2005) at structureHarvester.py v0.7 (Earl and VonHoldt 2012). The graphical output was constructed using the Clumpak (Kopelman et al. 2015). Analysis of molecular variance (AMOVA) was conducted by amova.poppr() function of package *poppr* to estimate the percent variation between and within the host, state of origin, mating type, mitochondrial haplotype, and genotype. The significance of estimated variance components was assessed based on 10,000 random permutations by function randtest() using *ade4* package (Dray and Dufour 2007). To validate the population clustering by the best K and the significant variation estimated, an analysis of principal components (PCA) and discriminant analysis of principal components (DAPC) was done by functions prcomp() and dapc() of *stats* (R Core Team 2024) and *adegenet* (Jombart 2008) packages, respectively. Minimum spanning networks (MSN) were generated to investigate clustering according to host, state of origin, mating type, mitochondrial haplotype, and genotype. Bruvo’s genetic distance was used as described before for MSN estimation.

The sequences of the four loci were manually inspected and the contig sequence was generated using SeqAssem (Hepperle 2004). The contigs were queried using BLAST in NCBI to confirm the similarity of gene and pathogen. The sequences were aligned in MEGA 11 (Tamura et al. 2021) with the available reference sequences (Coomber et al. 2023). The phylogenetic trees were inferred for each locus individually, for all loci concatenated, for loci *Btub*, *TigA*, and *ITS* concatenated, and for loci *Btub* and *TigA* concatenated by the Bayesian method.

Bayesian phylogenies were reconstructed under the best likelihood model for each locus, using MCMC with 5,000,000 repetitions in MrBayes on XSEDE (Ronquist et al. 2012). The statistical selection of the best-fit evolutionary model of nucleotide substitution for each locus was estimated using the Bayesian Information Criterion (BIC) at jModelTest2 on ACCESS (Darriba and Posada 2014). The phylogenetic trees were plotted on FigTree v1.4.4 (Rambaut et al. 2014) and exported to the graphic edition in CorelDRAW® 2021.

## Results

### Mating type characterization. PCR-based method

One hundred thirty-five isolates were characterized by mating type using PCR-based method and 52 were paired on culture medium. Both mating types, A1 (67 isolates) and A2 (68 isolates), were identified in the Brazilian population of *P*. *infestans* by molecular markers. The pairing on culture medium identified the mating types A1 (42 isolates) and A2 (13 isolates), two A1/A2 isolates, and zero SF (Table 1).

**Table 1.** Characterization of the isolates from the Brazilian population of *Phytophthora infestans* by geographic origin (Municipality), year of sample (Year), host of origin (Host), mating type by molecular markers and by pairing on culture medium, mitochondrial DNA haplotype, mefen<u>oxam sensitivity, and SSR genotype.</u>

| Isolate | Municipality | Year | Host | Mating type |  | Mitochondrial DNA | Mefenoxam Sensitivity* | SSR Genotype |
| --- | --- | --- | --- | --- | --- | --- | --- | --- |
|  |  |  |  | by molecular markers | by pairing |  |  |  |
| Pi1 | Viçosa - MG | 2020 | Potato | A2 | - | lb | - | SA_1_A2 |
| Pi2 | Patrocínio - MG | 2020 | Potato | A1 | A1 | la | - | EU_2_A1 |
| Pi3 | Perdizes - MG | 2020 | Potato | A1 | A1 | la | - | EU_2_A1 |
| Pi4 | Perdizes - MG | 2020 | Potato | A1 | A1 | la | - | EU_2_A1 |
| Pi5 | São Gotardo - MG | 2020 | Potato | A1 | A1 | la | - | EU_2_A1 |
| Pi6 | São Gotardo - MG | 2020 | Potato | A1 | A1 | la | - | EU_2_A1 |
| Pi7 | São Gotardo - MG | 2020 | Potato | A1 | A1 | la | - | EU_2_A1 |
| Pi8 | São Gotardo - MG | 2020 | Potato | A1 | A1 | la | - | EU_2_A1 |
| Pi9 | Rio Paranaíba - MG | 2020 | Potato | A1 | A1 | la | - | EU_2_A1 |
| Pi10 | São Gotardo - MG | 2020 | Potato | A1 | A1 | la | - | EU_2_A1 |
| Pi11 | São Gotardo - MG | 2020 | Potato | A1 | A1 | la | - | EU_2_A1 |
| Pi12 | São Gotardo - MG | 2020 | Potato | A1 | A1 | la | - | EU_2_A1 |
| Pi13 | São Gotardo - MG | 2020 | Potato | A1 | A1 | la | - | EU_2_A1 |
| Pi14 | São Gotardo - MG | 2020 | Potato | A1 | A1 | la | - | EU_2_A1 |
| Pi15 | São Gotardo - MG | 2020 | Potato | A1 | A1 | la | - | EU_2_A1 |
| Pi16 | São Gotardo - MG | 2020 | Potato | A1 | A1 | la | - | EU_2_A1 |
| Pi17 | São Gotardo - MG | 2020 | Potato | A1 | A1 | la | - | EU_2_A1 |
| Pi18 | São Gotardo - MG | 2020 | Potato | A1 | A1 | la | - | EU_2_A1 |
| Pi19 | São Gotardo - MG | 2020 | Potato | A1 | A1 | la | - | EU_2_A1 |
| Pi20 | Rio Paranaíba - MG | 2020 | Potato | A1 | A1 | la | - | EU_2_A1 |
| Pi21 | Rio Paranaíba - MG | 2020 | Potato | A1 | A1 | la | - | EU_2_A1 |
| Pi22 | Rio Paranaíba - MG | 2020 | Potato | A1 | A1 | la | - | EU_2_A1 |
| Pi23 | Rio Paranaíba - MG | 2020 | Potato | A1 | A1 | la | - | EU_2_A1 |
| Pi24 | Rio Paranaíba - MG | 2020 | Potato | A1 | A1 | la | - | EU_2_A1 |
| Pi25 | Rio Paranaíba - MG | 2020 | Potato | A1 | A1 | la | - | EU_2_A1 |
| Pi26 | Rio Paranaíba - MG | 2020 | Potato | A1 | A1 | la | - | EU_2_A1 |
| Pi27 | Rio Paranaíba - MG | 2020 | Potato | A1 | A1 | la | - | EU_2_A1 |
| Pi28 | Rio Paranaíba - MG | 2020 | Potato | A1 | A1 | la | - | EU_2_A1 |
| Pi29 | Rio Paranaíba - MG | 2020 | Potato | A1 | A1 | la | - | EU_2_A1 |
| Pi30 | Viçosa - MG | 2020 | Tomato | A2 | - | lb | - | SA_1_A2 |
| Pi31 | Rio Paranaíba - MG | 2020 | Potato | A1 | A1 | la | - | EU_2_A1 |
| Pi32 | Camanducaia - MG | 2020 | Potato | A2 | - | la | - | EU_37_A2 |
| Pi33 | Ipuiúna - MG | 2020 | Potato | A2 | - | la | - | EU_37_A2 |
| Pi34 | Espírito Santo do Dourado - MG | 2020 | Potato | A2 | - | la | - | EU_37_A2 |
| Pi35 | Camanducaia - MG | 2020 | Potato | A2 | - | la | - | EU_37_A2 |
| Pi36 | Camanducaia - MG | 2020 | Potato | A2 | - | la | - | EU_37_A2 |
| Pi37 | Camanducaia - MG | 2020 | Potato | A2 | - | la | - | EU_37_A2 |
| Pi38 | Camanducaia - MG | 2020 | Potato | A2 | A2 | la | S | EU_37_A2 |
| Pi39 | Ipuiúna - MG | 2020 | Potato | A2 | - | la | - | EU_37_A2 |
| Pi40 | Ipuiúna - MG | 2020 | Potato | A2 | A2 | la | R | EU_37_A2 |
| Pi41 | Espírito Santo do Dourado - MG | 2020 | Potato | A2 | - | la | - | EU_37_A2 |
| Pi42 | Camanducaia - MG | 2020 | Potato | A2 | - | la | - | EU_37_A2 |
| Pi43 | Ipuiúna - MG | 2020 | Potato | A2 | - | la | - | EU_37_A2 |
| Pi44 | Espírito Santo do Dourado - MG | 2020 | Potato | A2 | - | lb | - | EU_37_A2 |
| Pi45 | Espírito Santo do Dourado - MG | 2020 | Potato | A2 | A2 | lb | S | EU_37_A2 |
| Pi46 | Ipuiúna - MG | 2020 | Potato | A2 | - | lb | - | EU_37_A2 |
| Pi47 | Espírito Santo do Dourado - MG | 2020 | Potato | A2 | - | la | - | EU_37_A2 |
| Pi48 | Camanducaia - MG | 2020 | Potato | A2 | - | la | - | EU_37_A2 |
| Pi49 | São Francisco de Paula - RS | 2021 | Potato | A2 | - | la | - | EU_37_A2 |
| Pi50 | São Francisco de Paula - RS | 2021 | Potato | A2 | - | la | - | EU_37_A2 |
| Pi51 | São Francisco de Paula - RS | 2021 | Potato | A2 | - | la | - | EU_37_A2 |
| Pi52 | São Francisco de Paula - RS | 2021 | Potato | A2 | - | la | - | EU_37_A2 |
| Pi53 | Palmas - PR | 2021 | Potato | A2 | - | la | - | EU_37_A2 |
| Pi54 | São Francisco de Paula - RS | 2021 | Potato | A2 | - | la | - | EU_37_A2 |
| Pi55 | Guarapuava - PR | 2021 | Potato | A1 | A1 | la | I | EU_2_A1 |
| Pi56 | São Francisco de Paula - RS | 2021 | Potato | A2 | - | la | - | EU_37_A2 |
| Pi57 | São Francisco de Paula - RS | 2021 | Potato | A2 | - | la | - | EU_37_A2 |
| Pi58 | São Francisco de Paula - RS | 2021 | Potato | A1 | - | la | - | EU_2_A1 |
| Pi59 | Ipuiúna - MG | 2020 | Potato | A1 | - | la | - | EU_37_A2 |
| Pi60 | Água Doce - SC | 2021 | Potato | A1 | - | la | - | EU_2_A1 |
| Pi61 | Água Doce - SC | 2021 | Potato | A1 | - | la | - | EU_2_A1 |
| Pi62 | Guarapuava - PR | 2021 | Potato | A1 | A1 | la | - | EU_2_A1 |
| Pi63 | São Francisco de Paula - RS | 2021 | Potato | A2 | - | la | - | EU_37_A2 |
| Pi64 | São Francisco de Paula - RS | 2021 | Potato | A2 | - | la | - | EU_37_A2 |
| Pi65 | São Francisco de Paula - RS | 2021 | Potato | A2 | - | la | - | EU_37_A2 |
| Pi66 | Guarapuava - PR | 2021 | Potato | A1 | - | la | - | EU_2_A1 |
| Pi67 | São Francisco de Paula - RS | 2021 | Potato | A2 | - | la | S | EU_37_A2 |
| Pi68 | São Francisco de Paula - RS | 2021 | Potato | A2 | - | la | - | EU_37_A2 |
| Pi69 | São Francisco de Paula - RS | 2021 | Potato | A2 | - | la | - | EU_37_A2 |
| Pi70 | Guarapuava - PR | 2021 | Potato | A2 | A2 | la | I | EU_2_A1 |
| Pi71 | Palmas - PR | 2021 | Potato | A1 | - | la | - | EU_2_A1 |
| Pi72 | Palmas - PR | 2021 | Potato | A1 | - | lb | - | EU_2_A1 |
| Pi73 | Água Doce - SC | 2021 | Potato | A1 | - | la | I | EU_2_A1 |
| Pi74 | Palmas - PR | 2021 | Potato | A1 | - | la | I | EU_2_A1 |
| Pi75 | Calmon - SC | 2021 | Potato | A1 | - | la | - | EU_2_A1 |
| Pi76 | Guarapuava - PR | 2021 | Potato | A1 | - | la | I | EU_2_A1 |
| Pi77 | Guarapuava - PR | 2021 | Potato | A1 | - | la | - | EU_2_A1 |
| Pi78 | Ipuiúna - MG | 2020 | Potato | A2 | - | la | - | EU_37_A2 |
| Pi79 | Bom Repouso - MG | 2020 | Potato | A2 | - | la | - | EU_37_A2 |
| Pi80 | São José dos Ausentes - RS | 2021 | Potato | A1 | - | - | - | EU_2_A1 |
| Pi81 | Guarapuava - PR | 2021 | Potato | A1 | - | la | - | EU_2_A1 |
| Pi82 | Camanducaia - MG | 2020 | Potato | A2 | A2 | lb | S | EU_37_A2 |
| Pi83 | Camanducaia - MG | 2020 | Potato | A2 | A2 | la | S | EU_37_A2 |
| Pi84 | Camanducaia - MG | 2020 | Potato | A2 | - | la | - | EU_37_A2 |
| Pi85 | Palmas - PR | 2021 | Potato | A2 | - | - | - | EU_37_A2 |
| Pi86 | Guarapuava - PR | 2021 | Potato | A1 | - | la | R | EU_2_A1 |
| Pi87 | Guarapuava - PR | 2021 | Potato | A1 | - | la | - | EU_2_A1 |
| Pi88 | Água Doce - SC | 2021 | Potato | A1 | A1 | la | I | EU_2_A1 |
| Pi89 | Água Doce - SC | 2021 | Potato | A1 | - | la | - | EU_2_A1 |
| Pi90 | Tatuí - SP | 2021 | Potato | A1 | - | la | I | EU_2_A1 |
| Pi91 | Tatuí - SP | 2021 | Potato | A1 | - | la | - | EU_2_A1 |
| Pi92 | Tatuí - SP | 2021 | Potato | A1 | - | la | - | EU_2_A1 |
| Pi93 | Guarapuava - PR | 2021 | Potato | A1 | - | la | - | EU_2_A1 |
| Pi94 | Mucugê - BA | 2021 | Potato | A2 | A2 | la | S | EU_37_A2 |
| Pi95 | Itapetininga - SP | 2021 | Potato | A1 | A1 | la | I | EU_2_A1 |
| Pi96 | Mucugê - BA | 2021 | Potato | A2 | - | la | S | EU_37_A2 |
| Pi97 | Tatuí - SP | 2021 | Potato | A1 | A1 | la | I | EU_2_A1 |
| Pi98 | Tatuí - SP | 2021 | Potato | A1 | A1 | la | R | EU_2_A1 |
| Pi99 | Palmas - PR | 2021 | Potato | A2 | - | la | - | EU_37_A2 |
| Pi100 | Mucugê - BA | 2021 | Potato | A2 | A2 | la | - | EU_37_A2 |
| Pi101 | Tatuí - SP | 2021 | Potato | A1 | A1 | la | I | EU_2_A1 |
| Pi102 | Calmon - SC | 2021 | Potato | A1 | A1/A2 | la | S | EU_2_A1 |
| Pi103 | Mucugê - BA | 2021 | Potato | A2 | - | la | S | EU_37_A2 |
| Pi104 | Água Doce - SC | 2021 | Potato | A1 | - | la | I | EU_2_A1 |
| Pi105 | Água Doce - SC | 2021 | Potato | A1 | - | la | - | EU_2_A1 |
| Pi106 | Água Doce - SC | 2021 | Potato | A1 | - | la | - | EU_2_A1 |
| Pi107 | Mucugê - BA | 2021 | Potato | A2 | A2 | la | S | EU_37_A2 |
| Pi108 | Santa Maria de Jetibá - ES | 2021 | Tomato | A2 | - | lb | I | SA_1_A2 |
| Pi109 | Afonso Cláudio - ES | 2021 | Tomato | A2 | - | lb | - | SA_1_A2 |
| Pi110 | Afonso Cláudio - ES | 2021 | Tomato | A2 | - | lb | - | SA_1_A2 |
| Pi111 | Afonso Cláudio - ES | 2021 | Tomato | A2 | - | lb | - | SA_1_A2 |
| Pi112 | Coimbra - MG | 2021 | Tomato | A2 | - | lb | - | SA_1_A2 |
| Pi113 | Palmas - PR | 2021 | Potato | A2 | - | la | - | EU_37_A2 |
| Pi114 | Palmas - PR | 2021 | Potato | A1 | - | la | - | EU_2_A1 |
| Pi115 | São Francisco de Paula - RS | 2021 | Potato | A2 | - | la | - | EU_37_A2 |
| Pi116 | Água Doce - SC | 2021 | Potato | A1 | - | la | I | EU_2_A1 |
| Pi117 | Palmas - PR | 2021 | Potato | A1 | - | la | - | EU_2_A1 |
| Pi118 | Palmas - PR | 2021 | Potato | A2 | A2 | la | S | EU_37_A2 |
| Pi119 | Viçosa - MG | 2022 | Tomato | A2 | - | lb | R | SA_1_A2 |
| Pi120 | Viçosa - MG | 2022 | Tomato | A2 | - | lb | - | SA_1_A2 |
| Pi121 | Palmas - PR | 2021 | Potato | A1 | A1 | la | I | EU_2_A1 |
| Pi122 | São Francisco de Paula - RS | 2022 | Potato | A1 | - | la | - | EU_2_A1 |
| Pi123 | Viçosa - MG | 2023 | Tomato | A2 | A2 | lb | - | SA_1_A2 |
| Pi124 | Espírito Santo do Dourado - MG | 2022 | Potato | A2 | A2 | la | S | EU_37_A2 |
| Pi125 | Espírito Santo do Dourado - MG | 2022 | Potato | A2 | A2 | la | R | EU_37_A2 |
| Pi126 | Calmon - SC | 2022 | Potato | A1 | A1/A2 | la | - | EU_2_A1 |
| Pi128 | Viçosa - MG | 2023 | Tomato | A2 | A1 | lb | I | SA_1_A2 |
| Pi132 | Domingos Martins - ES | 2024 | Tomato | - | A1 | - | - | - |
| Pi133 | Castelo - ES | 2024 | Tomato | A2 | - | lb | - | SA_1_A2 |
| Pi134 | Santa Maria de Jetibá - ES | 2024 | Tomato | A2 | - | lb | - | SA_1_A2 |
| Pi135 | Santa Maria de Jetibá - ES | 2024 | Tomato | A2 | A1 | lb | - | SA_1_A2 |
| Pi136 | Santa Maria de Jetibá - ES | 2024 | Tomato | A2 | A1 | lb | - | SA_1_A2 |
| Pi137 | Santa Maria de Jetibá - ES | 2024 | Tomato | A2 | A1 | lb | - | SA_1_A2 |
| Pi138 | Castelo - ES | 2024 | Tomato | A2 | - | lb | - | SA_1_A2 |

\*S = sensitive; I = intermediate; and R = resistant to mefenoxam.

After enzymatic digestion of the PCR product, 67 isolates presented amplification fragments of the A1 type (∼ 100, 500, and 600 bp) and 68 isolates of the A2 type (∼ 100 and 500 bp). The proportion of mating type (A1:A2) according to the state of origin was: 0:5 in BA, 30:31 in MG, 0:10 in ES, 7:0 in SP, 15:6 in PR, 12:0 in SC, and 3:14 in RS state (Figure 1A). Both mating types were identified co-occurring in the same fields in MG and PR states.

**Figure 1.**
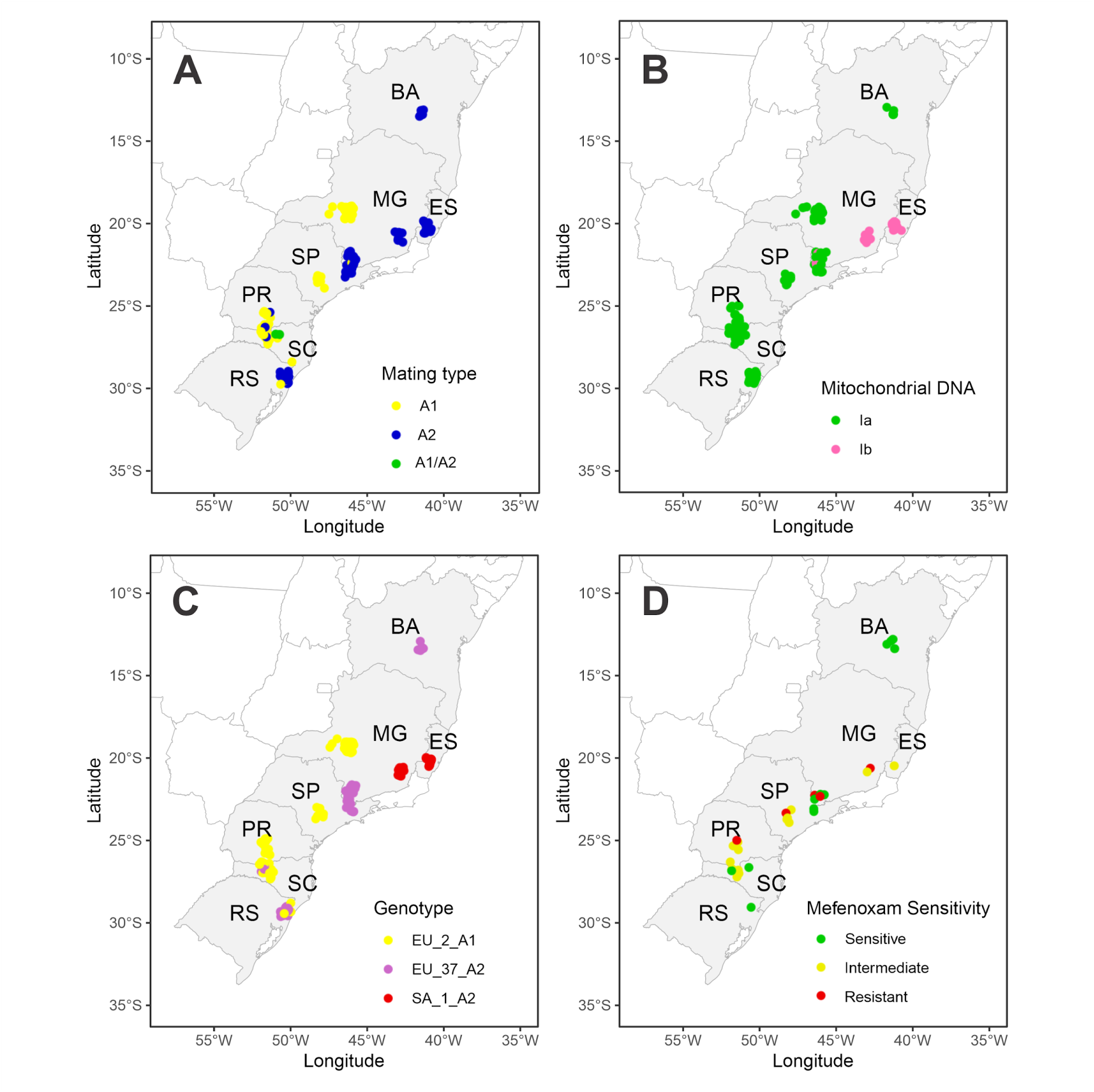
Isolate sites (dots) according to the state of origin: BA = Bahia, MG = Minas Gerais, SP = São Paulo, PR = Paraná, SC = Santa Catarina, and RS = Rio Grande do Sul. The color dots are according to phenotypic or genotypic characteristics as indicated on legends. A: mating type; B: Mitochondrial DNA haplotype; C: Genotype; and D. Mefenoxam sensitivity.

**Figure 1.**
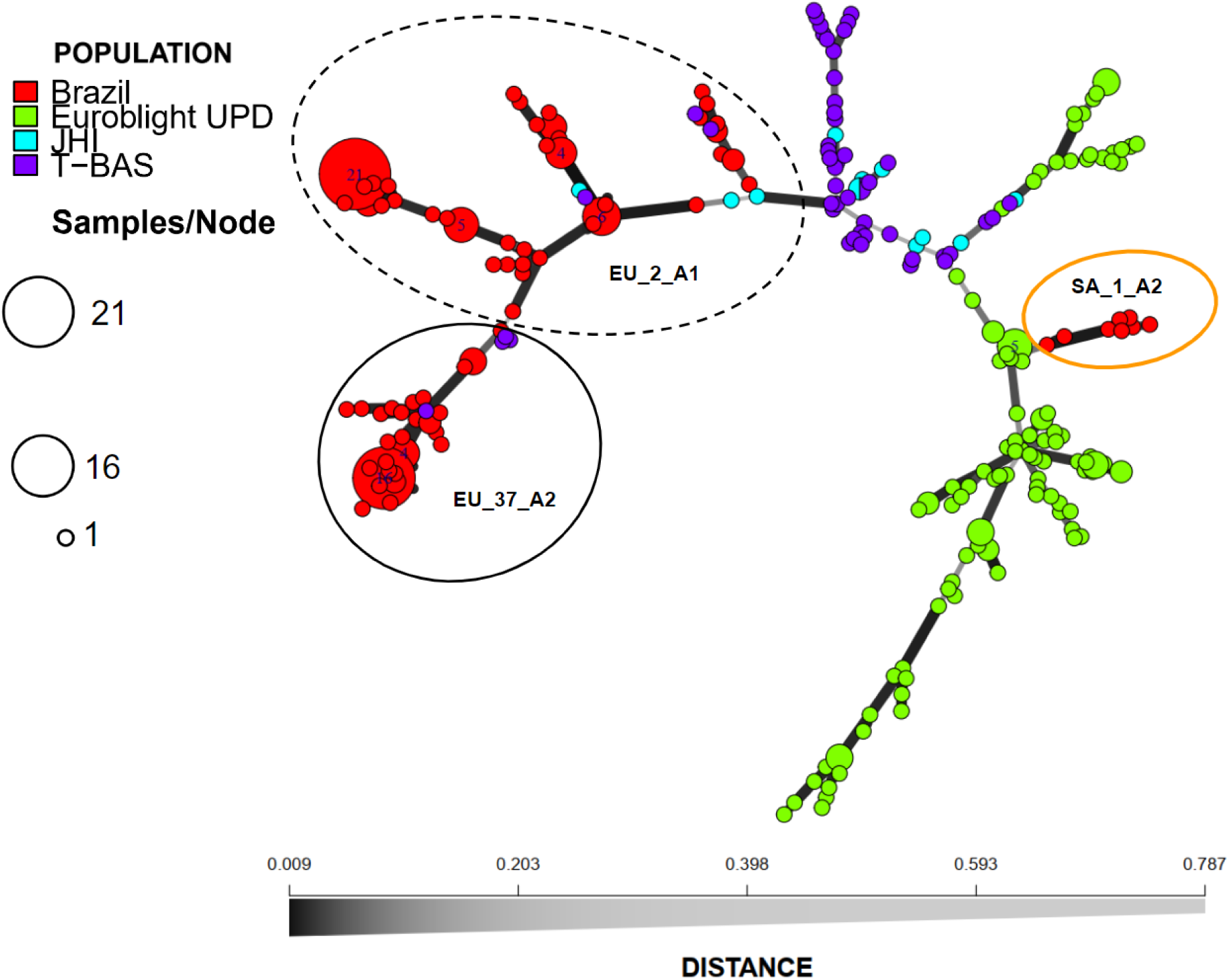
Minimum spanning network of genotypes of *Phytophthora infestans* based on 12-plex simple-sequence repeats (SSRs) using Bruvo’s distance. Nodes are colored by the data set: red, green, blue, and purple indicate the isolates from current population from Brazil, isolates from dataset of Saville and Ristaino (2019) study using methods in line with the Euroblight scoring system (Euroblight UPD), control isolates sent by Dr. David Cooke (JHI), and isolates from the dataset of Coomber et al. (2023) and lineages described in *P*. *infestans* classifier (T-BAS). The nodes represent the multilocus genotypes (MLGs), and their size is proportional to the number of isolates sharing a given MLG. The thickness and darkness of branches represent Bruvo’s genetic distance between adjacent nodes. The circle with a black dotted line grouped the isolates from the EU_2_A1 genotype, the circle with a black continuous line grouped isolates from the EU_37_A2 genotype, and the circle with an orange continuous line grouped the isolates from the SA_1_A2 genotype.

#### Pairing-method

Forty-two isolates formed oospores in the pairing assays on culture medium with the A2 tester, thus these isolates were characterized as A1 mating type; 13 were identified as A2 mating type because oospores were formed when paired with the A1 tester, and two isolates formed oospores with both A1 and A2 testers and were characterized as A1/A2. Most isolates identified by pairing as A1 and A1/A2 were identified as A1 by PCR. Five isolates from tomato were identified as A1 by pairing and as A2 by the PCR-based method, and all isolates identified as A2 by pairing culture were also identified as A2 by the PCR-based method. The proportion of mating type identified by the pairings in culture medium according to the state of origin was the same as described by the PCR-based method except for the ES and SC states. Isolates identified as SF were sampled from the SC state. The proportion of mating type (A1:A2:A1/A2) was 1:14:2 to SC state. Regarding the host of origin, the isolates from potato were identified as A1, A2, and A1/A2, and isolates from tomato were identified as A1 or A2.

#### Mitochondrial haplotypes

The mitochondrial haplotypes of 134 isolates were characterized. The haplotypes Ia (n = 110) and Ib (n = 24) were detected (Table 1). The distribution of mitochondrial haplotypes (Ia:Ib) according to the state of origin was: 5:0 in BA, 50:11 in MG, 0:10 in ES, 0:7 in SP, 20:1 in PR, 12:0 in SC, and 17:0 in RS (Figure 1B). The isolates from potato were identified as Ia and Ib, and the ones from the tomato were identified as Ib haplotype.

#### Mefenoxam sensitivity

Thirty-two isolates, 27 from potato and five from tomato, were assessed for sensitivity to mefenoxam. Fourteen isolates (12 from potato and two from tomato) were sensitive to mefenoxam, 14 from potato were classified as intermediate, and five isolates (two from potato and three from tomato) were resistant (Table 1).

The sensitive isolates were obtained from BA (n = 2), MG (n = 5), ES (n = 2), PR (n = 1), SC (n = 1), and RS (n = 1) states (Figure 1D). The intermediate isolates are from MG (n = 1), SP (n = 3), PR (n = 5), and SC (n = 5) (Figure 1D). The resistant isolates are from MG (n = 1), ES (n = 3), and SP (n = 1) (Figure 1D).

#### SSR genotyping

One hundred thirty-five isolates obtained from potato and tomato-producing areas sampled in the Brazilian states between the 2020 and 2024 seasons were genotyped by 12-plex SSR. Using a cut-off value of 0.128, 67 individuals were identified as EU_2_A1 genotype, 49 as EU_37_A2, and 19 did not match any genotype ever described and were named as SA_1_A2 genotype (Table 1). The MSN shows isolates from Brazil grouping with isolates previously identified as EU_2_A1 and EU_37_A2 genotype and some isolates not grouping with any known genotype (Figure 2). The EU_2_A1 isolates were found in MG, SP, PR, SC, and RS states, the EU_37_A2 were from BA, MG, PR, and RS states, and the SA_1_A2 were present in MG and ES states (Figure 1C).

**Figure 2.**
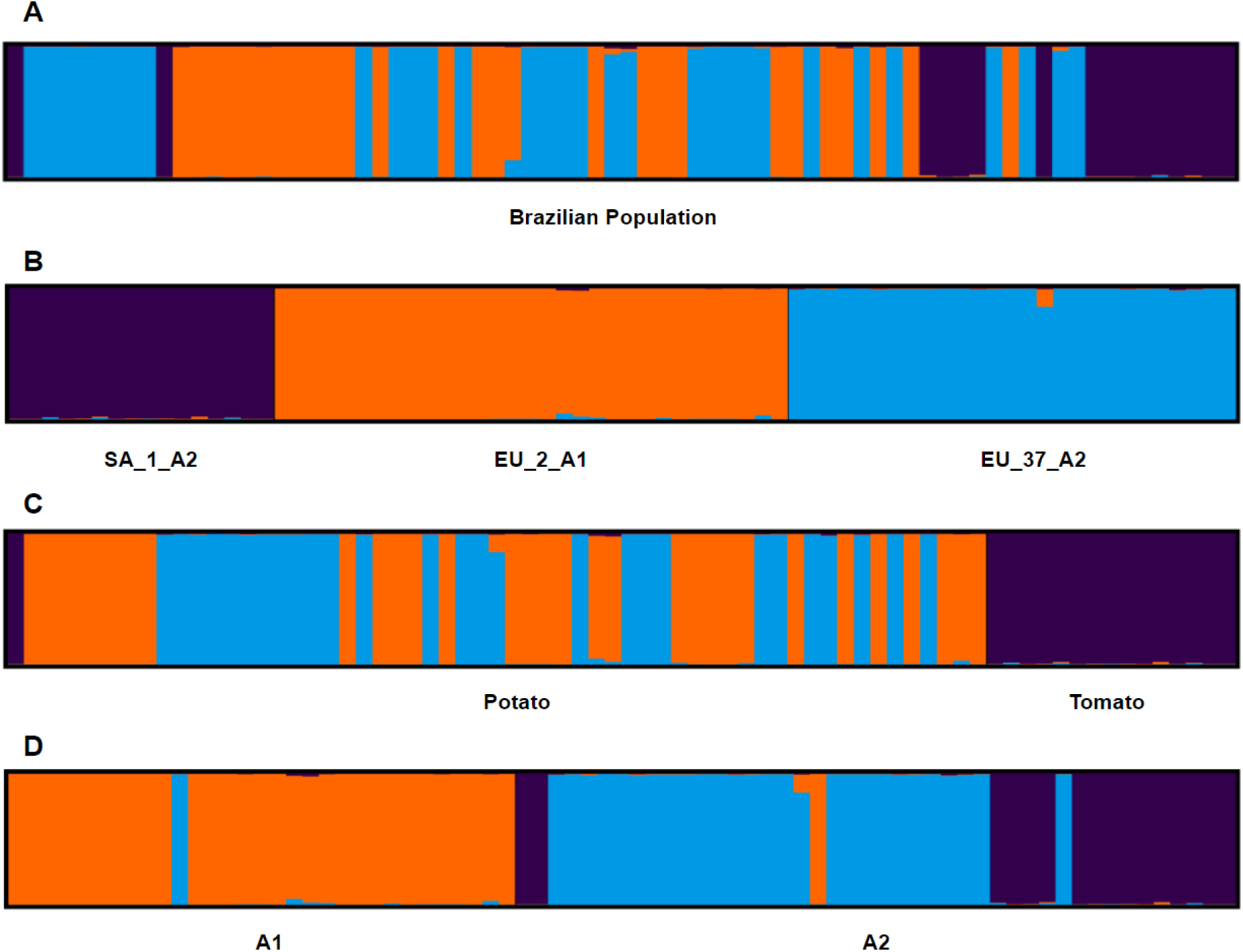
Cluster analysis of the current Brazilian population of *Phytophthora infestans*. The analysis was done considering the entire population without phenotype information (A); the genotypes identified (B); the host of origin (C); and the mating type characterized by molecular markers. The population was analyzed assuming three genetic groups (K = 3). Each bar in the figures represents a multilocus genotype (MLG) colored by the proportion of each MLG variation that belongs to one of the three K groups.

#### Population variability

In the entire population, 63 alleles were observed. The average number of alleles was 5.25. The D13 locus had the highest number of alleles (Allele num = 12), and the SSR11 and SSR2 loci had the lowest allele number (Allele num = 2) (Table 2). The SSR4 locus has the highest diversity according to the Simpson index (1-D = 0.82), and the SSR11 locus had the highest evenness (Evenness = 1.00) (Table 2). The percentage of missing data was 0.19 %. The observed heterozygosity was 1 for all loci, and the expected heterozygosity varied between 0.50 (SSR2) to 0.85 (SSR4) with 0.69 of the mean (Table 2).

**Table 2.**
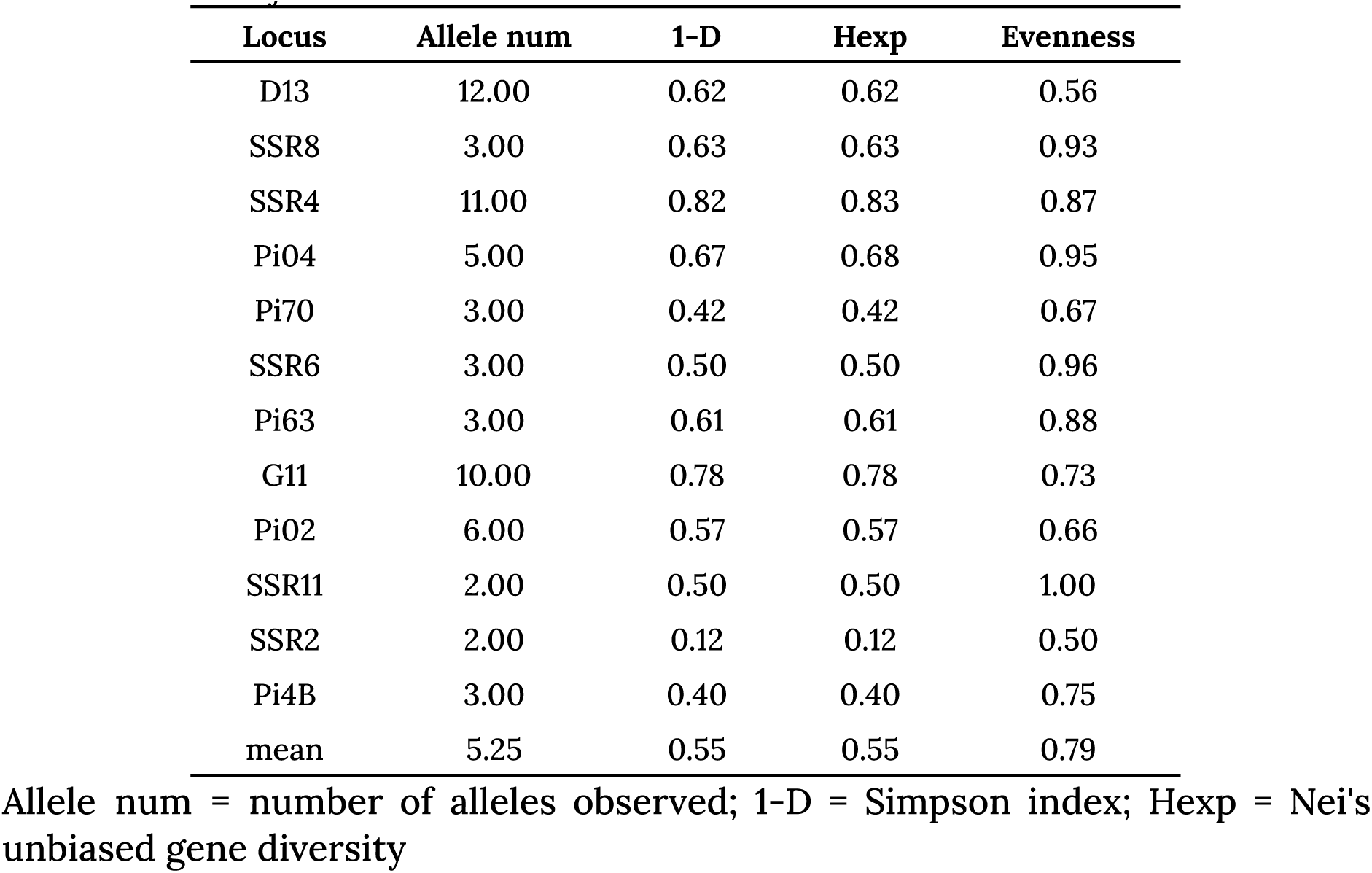
Population statistics by locus observed in the Brazilian population of *Phytophthora infestans*.

A total of 74 original multilocus genotypes (MLGs) were observed across 135 triploid individuals based on 12 codominant loci. Thirteen MLGs occurred in more than one sampled individual. The genotype EU_2_A1 had a higher number of MLGs than the other genotypes presented in the population and was the most diverse according to the Shannon-Wiener index (H = 2.79) (Table 3). However, the genotype SA_1_A2 had the highest evenness statistic (E.5 = 0.89) (Table 3). The potato host presented high MLG diversity (H = 3.47), but low evenness (E.5 = 0.47) (Table 2). The subpopulation of RS state was the most diverse (H = 2.64), and those of BA and ES had the highest evenness (H = 1 and 0.96, respectively) (Table 3). According to the index of association, asexual reproduction is predominant in the Brazilian population in all subpopulation assignments (host, state, etc.) (Ia and r̄d ≠ 0).

**Table 3.**
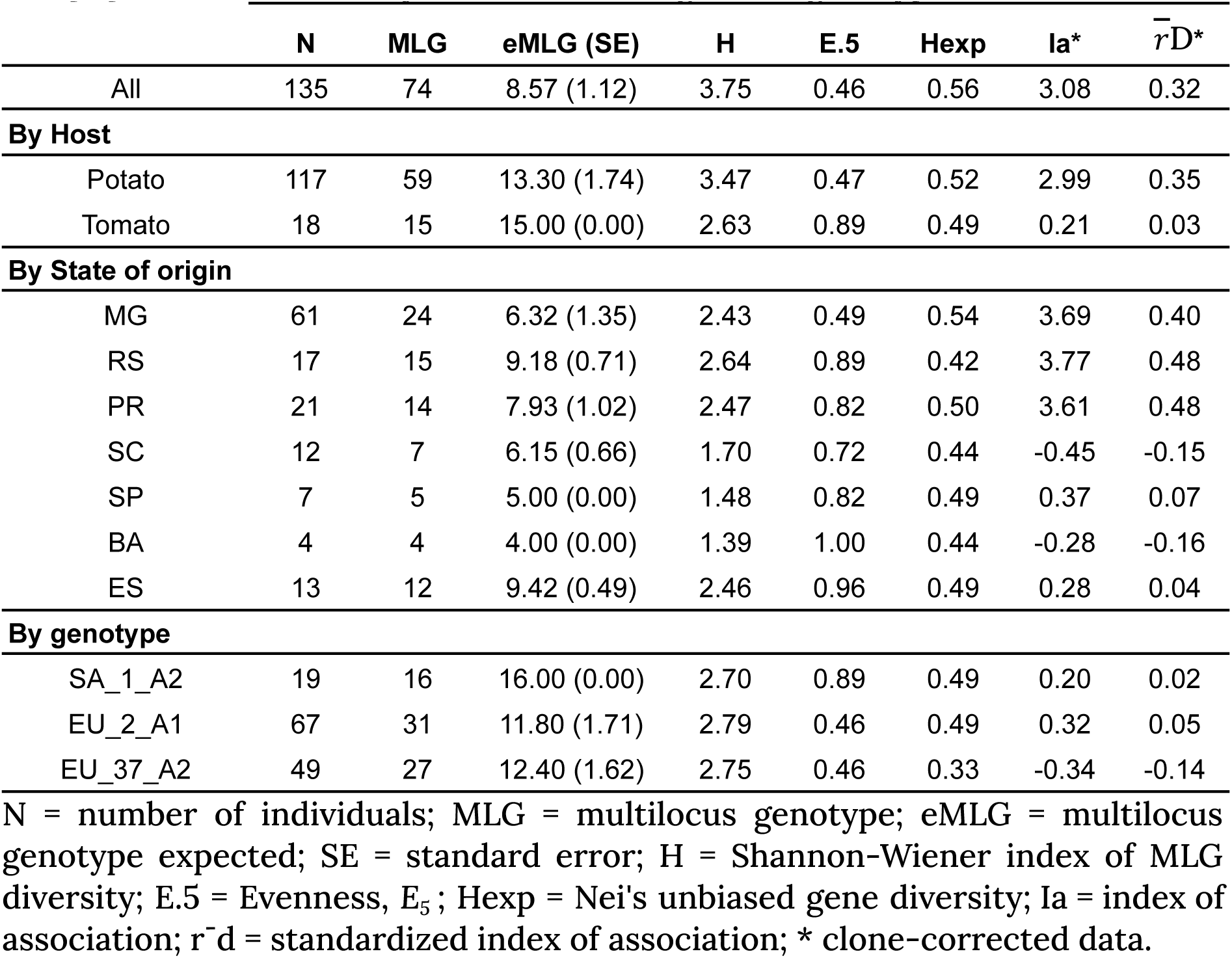
Population summary statistics for *Phytophthora infestans* from Brazil. The population is sorted by host, state of origin, and genotype.

No loci were at HWE across all 135 individuals. There was evidence to reject the null hypothesis of HWE when analyzing both, the original population data (with clones) and the clone-corrected dataset, and conclude that the population is on linkage disequilibrium (LD) (P < 0.001 and 𝑟D=0.418), a characteristic observed in clonal populations.

#### Population structure

The Brazilian population of *P*. *infestans* is structured into three clusters (Figure 2). The MSN also grouped the isolates into three clusters of genotypes (Figure 3A) which in turn were associated with hosts (Figure 3B). Two clusters were defined by host plants. The red group is composed of isolates sampled from tomato plants, except one that was obtained from potato plants cultivated in an experimental plot that was set in a tomato producing area (Viçosa, MG). The other two group comprised isolates sampled from potato plants (Figure 3B).

**Figure 3.**
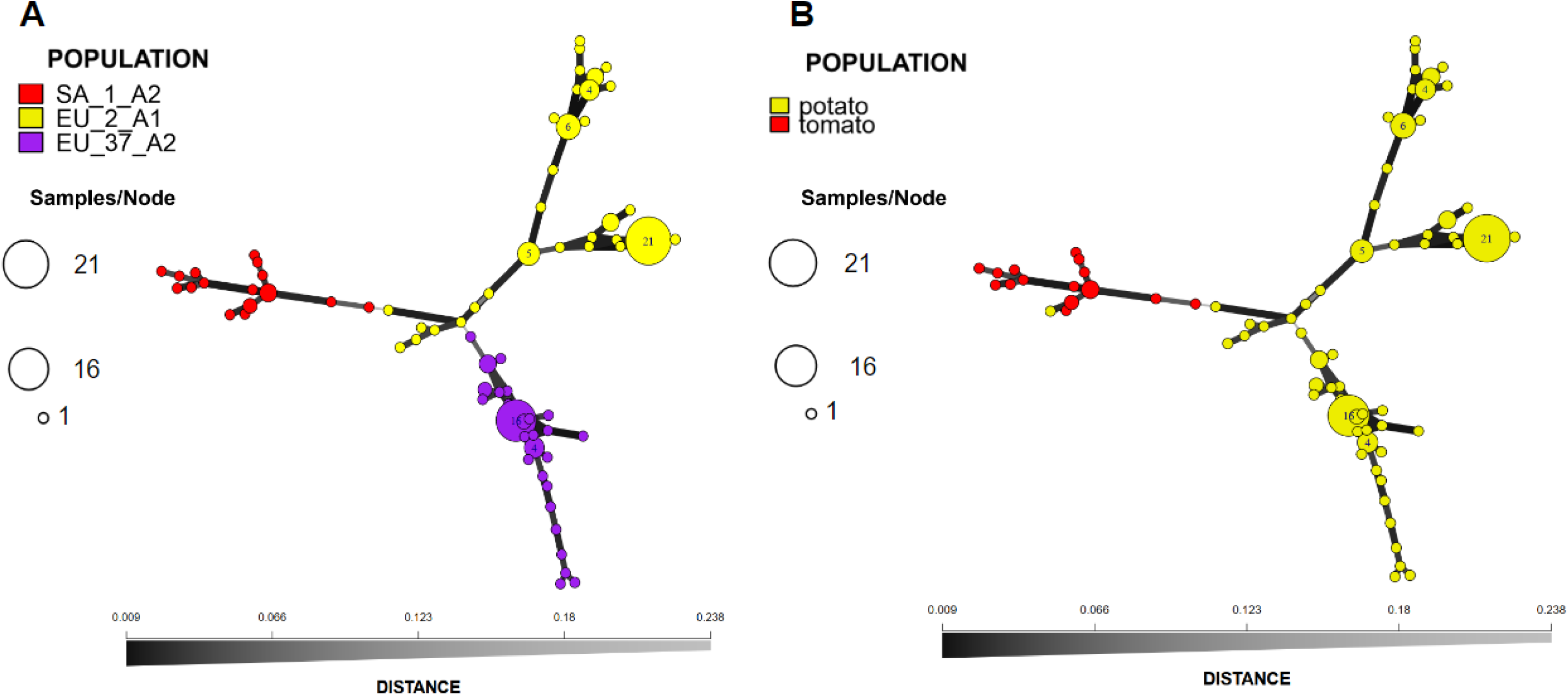
Minimum spanning network of Brazilian population of *Phytophthora infestans* based on 12-plex simple-sequence repeats (SSRs) using Bruvo’s distance. Nodes are coloured by genotypes (A) or host (B), represent the multilocus genotypes (MLGs), and their sizes increase according to the number of isolates sharing the MLGs. The thickness and darkness of branches represent the Bruvo’s genetic distance between adjacent nodes.

The NJ tree constructed with Bruvo’s distance grouped the isolates into three well supported clades (bootstrap > 70%) (Figure 4).

**Figure 4.**
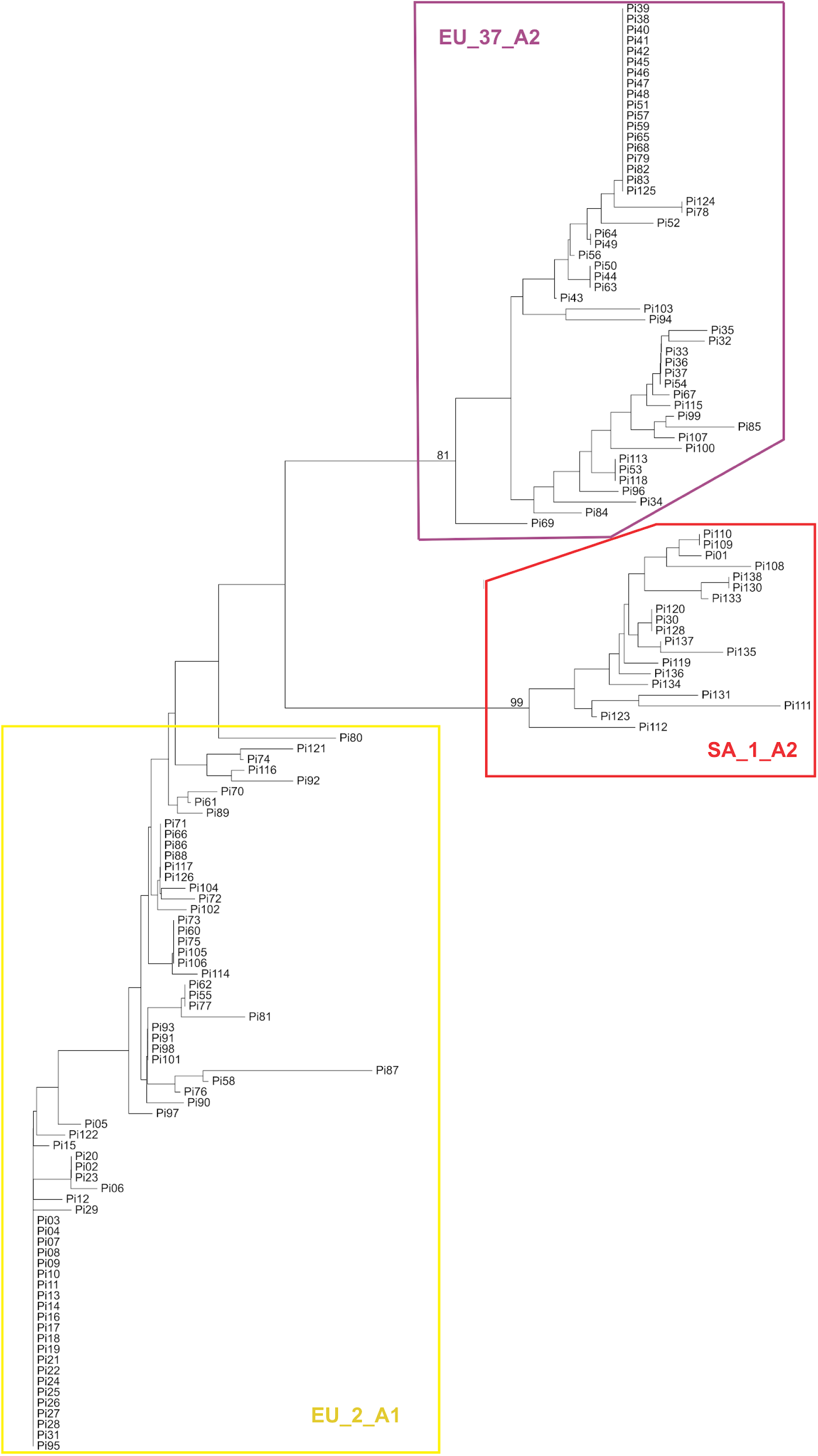
Neighbor-joining tree of Brazilian population of *Phytophthora infestans* sampled between 2020 and 2024, based on 12-plex simple-sequence repeats (SSRs) using Bruvo’s genetic distance. The isolates are color labeled according to the genotype. Purple: EU_37_A2 genotype; Red: SA_1_A2 genotype; and yellow: EU_2_A1 genotype.

The AMOVA results indicate that there was significant variation between genotype (75.7 %, p < 0.001), and host (51.2 %, p < 0.001). The PCA and DAPC support the population clustering according to the SSR genotypes (Figure 5).

**Figure 5.**
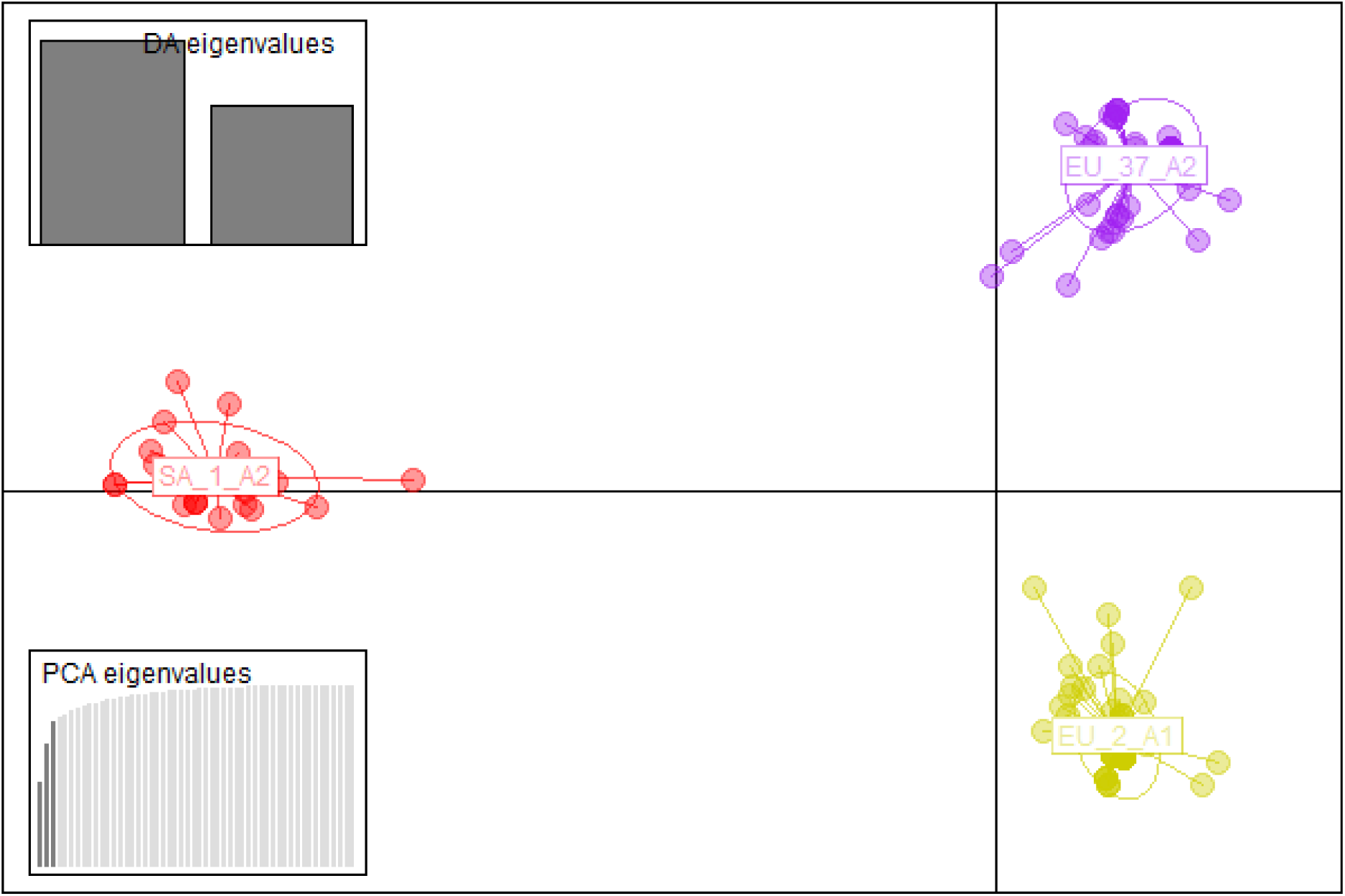
Discriminant analysis of principal components of the Brazilian population of *Phytophthora infestans* sampled between 2020 and 2024. The isolates were grouped by their respective genotypes.

Temporal analysis. A total of 170 MLGs were observed across 301 triploid individuals from 1998 to 2024 by six codominant loci. The 1998-2010 subpopulation had the highest diversity and evenness according to the Shannon-Wiener index and E.5 index (H = 4.58; E.5 = 0.70) (Table 4). There is evidence of the probable occurrence of sexual reproduction in the 1998-2010 population (Ia = 0.09; r̄d = 0.02) (Table 4).

**Table 4.** Population summary statistics for *Phytophthora infestans* from Brazil. Subpopulation defined by year.

|  | N | MLG | eMLG (SE) | H | E.5 | Hexp | Ia* | r<sup>-</sup>d* |
| --- | --- | --- | --- | --- | --- | --- | --- | --- |
| All | 301 | 170 | 89.70 (4.02) | 4.64 | 0.45 | 0.65 | 0.77 | 0.155 |
| **2020-2024** | 135 | 50 | 50.00 (0.00) | 3.17 | 0.48 | 0.65 | 1.83 | 0.37 |
| **1998-2010** | 166 | 120 | 101.20 (2.36) | 4.58 | 0.70 | 0.50 | 0.09 | 0.02 |

N = number of individuals; MLG = multilocus genotype; eMLG = multilocus genotype expected; SE = standard error; H = Shannon-Wiener index of MLG diversity; E.5 = Evenness, E<sub>5</sub>; Hexp = Nei's unbiased gene diversity; Ia = index of association; r<sup>-</sup>d = standardized index of association; \* clone corrected value.

The mean number of observed alleles per locus for the 1998 to 2024 subpopulation was 11.5 (Table 5). The total number of alleles was 58 and 37 for the 2020-2024 and 1998-2010 population, respectively. The diversity of locus G11 was the highest (1-D = 0.83) and the Pi04 locus had highest evenness (Evenness = 0.64) (Table 5). The percentage of missing data was 0.17 %. The observed heterozygosity was 1 in both populations (2020-2024 and 1998-2010) and the heterozygosity expected ranged between 0.66 (Pi70) to 0.83 (G11) to 2020-2024 population and from 0.53 (Pi02) to 0.79 (G11) to 1998-2010 population (Table 5).

**Table 5.**
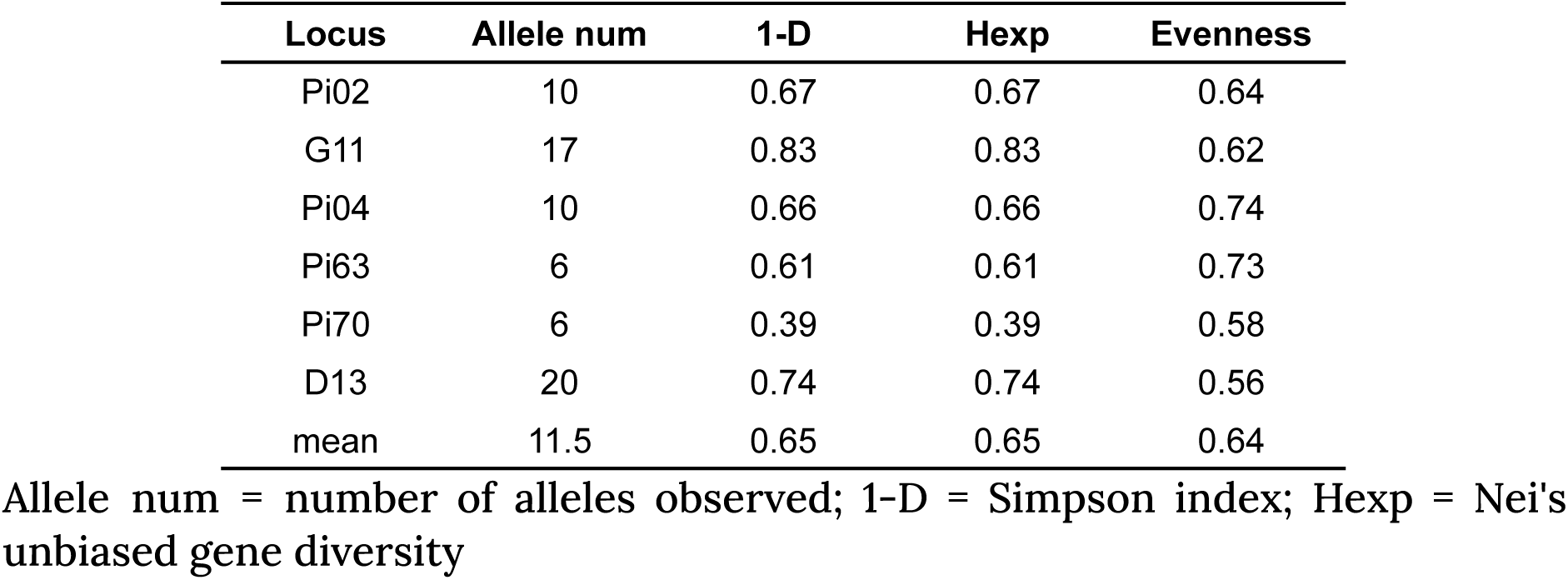
Population statistics of 2020-2024 and 1998-2010 populations, by locus observed in the Brazilian population of *Phytophthora infestans*.

| Locus | Allele num | 1-D | Hexp | Evenness |
| --- | --- | --- | --- | --- |
| Pi02 | 10 | 0.67 | 0.67 | 0.64 |
| G11 | 17 | 0.83 | 0.83 | 0.62 |
| Pi04 | 10 | 0.66 | 0.66 | 0.74 |
| Pi63 | 6 | 0.61 | 0.61 | 0.73 |
| Pi70 | 6 | 0.39 | 0.39 | 0.58 |
| D13 | 20 | 0.74 | 0.74 | 0.56 |
| mean | 11.5 | 0.65 | 0.65 | 0.64 |

Allele num = number of alleles observed; 1-D = Simpson index; Hexp = Nei's unbiased gene diversity

The number of genetic clusters varied according to the sampling date of the isolates. A single cluster describes the past (1998-2010) subpopulation, while two clusters were inferred for the current (2020-2024) subpopulation (Figure 6).

**Figure 6.**
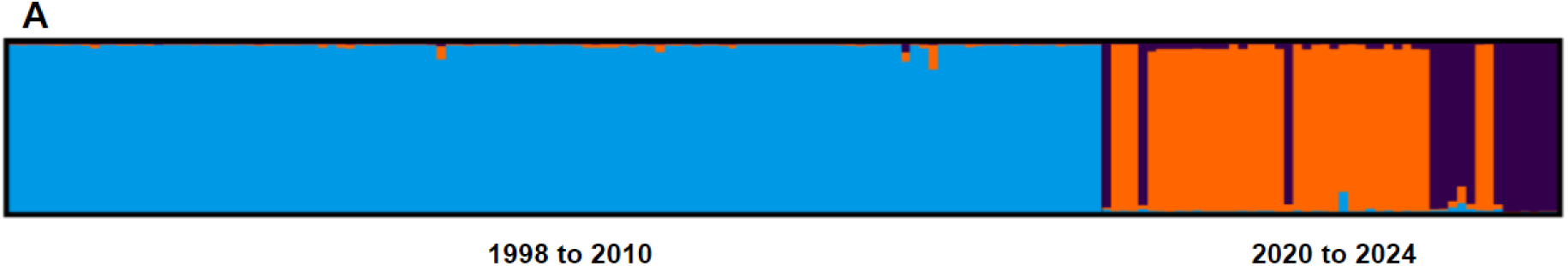
Bayesian 50 % majority rule consensus tree based on the *Btub*, G3PDH, and TIG genes of 27 Brazilian isolates of *Phytophthora infestans* obtained from potato (n = 11) and tomato (n = 16) and reference sequences retrieved from GenBank (isolate description shown in the tree). Bayesian posterior probabilities were also calculated and values are shown at each node > 0.80. The tree was rooted in *P*. *quercina* (Clade 3). The scale bar represents the expected number of substitutions per site. The Brazilian isolates used in the analyses are indicated by their code in bold. The genotypes and host of origin of isolates from Brazil are indicated by a color bar and the respective genotype or host. Red, purple, and yellow bars adjacent to the node labels followed by SA_1_A2, EU_37_A2, or EU_2_A1 indicate these genotypes in that order. Yellow and red bars on the right side followed by Potato or Tomato indicate the hosts of origin.

**Figure 6.**
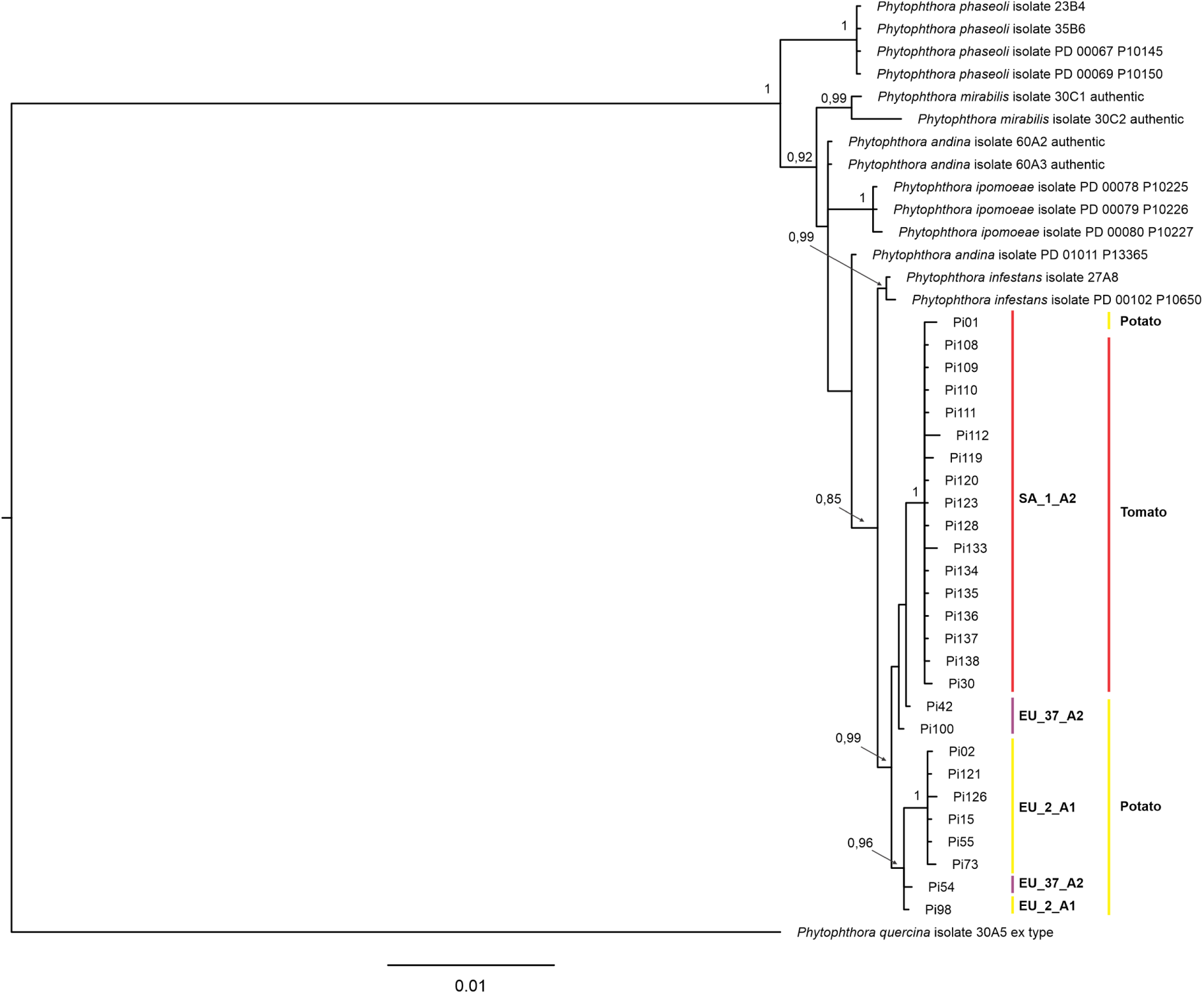
Cluster analysis of the past (1998 to 2010) and current (2020 to 2024) Brazilian population of *Phytophthora infestans*. The population was analyzed assuming three (A) or five (B) genetic groups (K = 3 or K = 5). Each bar in the figures represents a multilocus genotype (MLG) defined based on SSR markers colored by the proportion of each MLG variation that belongs to one of the three or five K groups.

The haplotypes from the past and current subpopulations formed two clusters in the MSN (Figure 7). The AMOVA results indicate significant variation within samples (65.60 %, P < 0.001).

**Figure 7.**
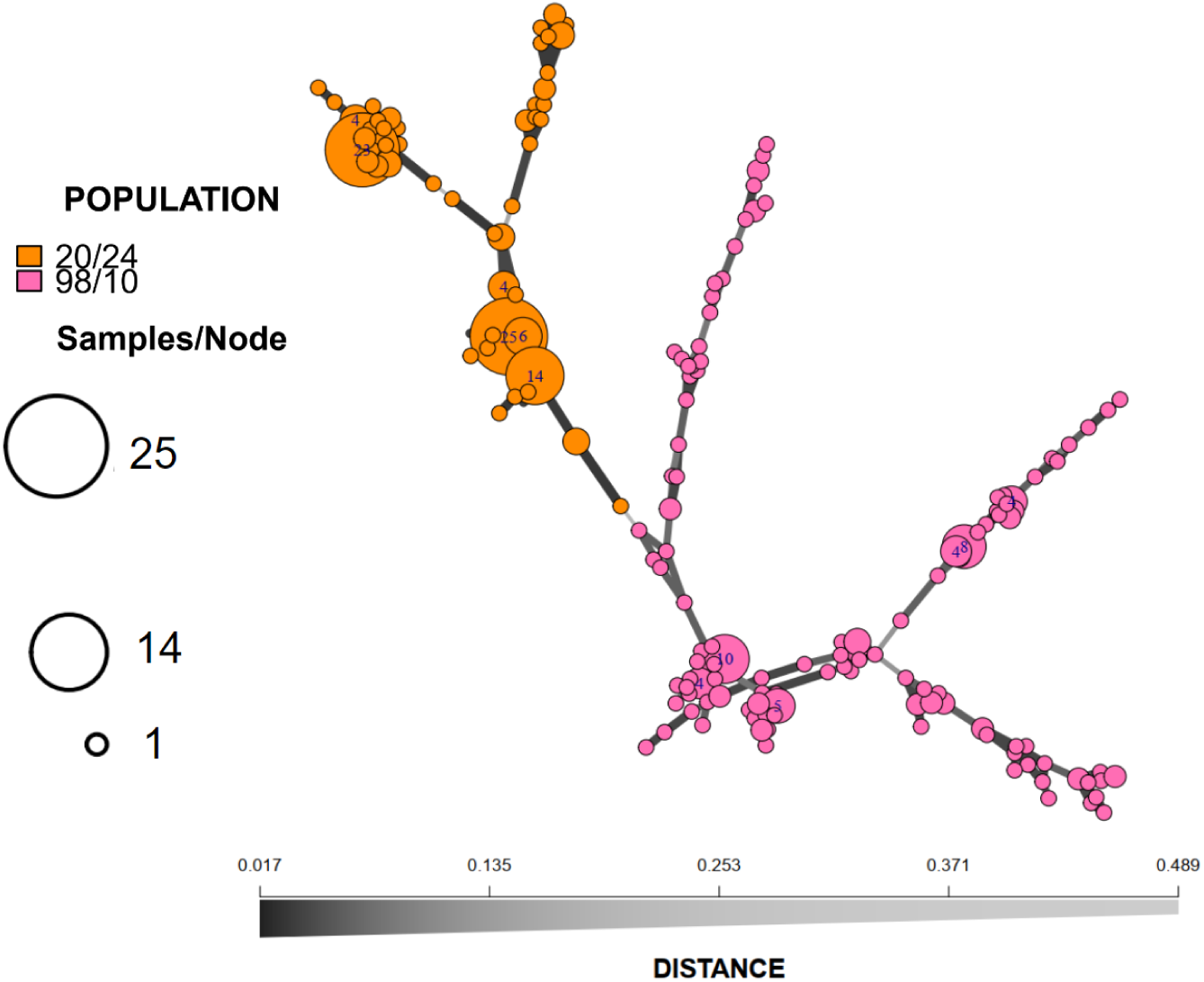
Minimum spanning network of Brazilian population of *Phytophthora infestans* sampled from 1998 to 2010 (pink) and from 2020 to 2024 (orange) based on 6 simple-sequence repeats (SSRs) using Bruvo’s distance. Nodes are colored by genotypes and represent the multilocus genotypes (MLGs) defined according to SSR markers, and their size increases according to the number of isolates sharing the MLGs. The thickness and darkness of branches represent Bruvo’s genetic distance between adjacent nodes.

#### Phylogenetic study

Partial sequences of the *Btub*, *TigA*, *ITS*, and *CoxI* loci were obtained for 27 isolates: 11 from potato and 16 from tomato. The best-fit evolutionary model chosen was GTR+G (nts = 6; rates = gamma) for *Btub*, GTR+I+G (nts = 6; rates = invgamma) for G3PDH (*TigA* loci), GTR+I (nts = 6; rates = propinv) for Tig (*TigA* loci) and *CoxI*, and HKY+I (nts = 2 rates = propinv) for *ITS* loci.

The aligned sequences from *Btub* (987 aligned characters), G3PDH (527 aligned characters), TIG (576 aligned characters), *ITS* (726 aligned characters), and *CoxI* (648 aligned characters) resulted in 94, 73, 86, 71, and 96 single nucleotide polymorphisms, respectively.

Initially, the phylogenetic analysis was done with all species of *Phytophthora* Clade 1. As the Brazilian isolates grouped in Clade 1c in all analyses, we proceed with the analysis only with species of this clade. The phylogenetic analysis of *ITS* and *CoxI* were able to separate the *Phytophthora* species among their clades but were not able to differentiate the species inside clade 1c (Figure S2, and S3). To compare potato and tomato isolates, the *ITS* and *CoxI* sequences were not used in the multigene analysis.

The multigene analysis was done with *Btub* and *TigA* loci. The alignment for the three concatenated genes (*Btub*, G3PDH, and TIG) contained 2,635 characters, including alignment gaps. The Bayesian phylogeny of the three genes distinguished the species: *P*. *phaseoli*, *P*. *mirabilis*, *P*. *andina*, *P*. *ipomoeae*, and *P*. *infestans*. The *P*. *infestans* isolates from Brazil of SA_1_A2 genotype, EU_2_A1 and EU_37_A2 genotypes grouped separately (Figure 6).

## Discussion

The genotypes of the current Brazilian population of *P*. *infestans* are different from the previous ones. Starting from the analysis of mating types, the presence of A1, A2, and A1/A2 isolates already indicate major changes in the population of the pathogen. Consequently, shifts in other markers were also noteworthy: Ia and Ib mitochondrial DNA haplotypes, and three clonal lineages EU_2_A1, EU_37_A2, and SA_1_A2, the new lineage, co-occurring in different potato and tomato producing regions. On the other hand, using sensitivity to mefenoxam as a “marker”, as formerly used in past studies of characterization of *P*. *infestans* populations, did not indicate any change in the population; i.e., populations of *P*. *infestans* still harbor sensitive, intermediate, and resistant individuals. In addition, supporting the phenotype differences observed in isolates from both hosts, Bayesian phylogeny distinguished two host-defined populations: one associated with potato and another with tomato. This study provides an update on the *P*. *infestans* population in Brazil, reported the A2 mating type in tomato fields for the first time, and described a new genotype associated with tomato.

The current Brazilian population of *P*. *infestans* has a new configuration. Until now, individuals of A1 mating type were almost exclusively sampled from tomato fields (Brommonschenkel 1988; Goodwin et al. 1994; Oliveira 2010; Reis et al. 2003, 2006; Santana et al. 2013; Zanotta 2019). In this study, all isolates obtained from tomato samples were characterized as A2 mating type. Thus, this is the first report of A2 mating type individuals occurring in tomato fields in Brazil. Furthermore, this signals a major shift in the population of *P*. *infestans* that causes late blight on tomato crops. Despite the genetic implications of this finding, its epidemiological impacts need to be better assessed because tomato isolates were obtained from two states only.

For a long time, the mating type most commonly reported in isolates obtained from potato fields was the A2 (Brommonschenkel 1988; Goodwin et al. 1994; Oliveira 2010; Reis et al. 2003, 2006; Santana et al. 2013) but recently, the mating type A1 was reported as dominant in potato fields (Zanotta 2019). The present study shows both mating types occurring at similar ratios on this host. These results can be explained by the current two genotypes occurring in potato fields in Brazil, EU_2_A1 and EU_37_A2 in very similar ratios.

The mating type characterization done by the PCR-based method was supported by the traditional pairing method in culture medium. The set of primers W16-1/W16-2 agreed 90.38 % with the results of pairing. The disagreement between methods occurred for five isolates obtained from tomato, and in isolates from potato identified as A1/A2 due to the formation of oospores with both testers. The first disagreement can be explained by the lack of consistency observed when pairing isolates from tomato among replicates (inconsistencies), probably due to large variation in time for isolates to grow, i.e. different mycelial growth rates. The second disagreement must be caused because the set of primers used was not able to identify A1/A2 mating type.

The oligonucleotides more commonly used are W16-1/W16-2 (Judelson et al. 1995), S1-A/S1-B (Judelson 1996), and PHYB-1/PHYB-2 (Kim and Lee 2002). These oligonucleotides allow the identification of both A1 and A2 mating types, respectively. Considering that efficient molecular markers for mating-types identification should present minimal error rates compared with pairing tests, primer validation is necessary due to the genetic variation of each population (Brylińska et al. 2018). The use and validation of molecular markers to identify the mating type is a practice that has been used (Beketova et al. 2014; Brylińska et al. 2018; Chowdappa et al. 2015; Dangi et al. 2021; Mazáková et al. 2010; Wharton et al. 2023; Zhang et al. 2006).

The Brazilian population was assessed for mating type using the three primer pairs (Zanotta 2019). In the present study, only the set of primers W16-1/W16-2 (Judelson et al. 1995) were useful, i.e. were reliable and agreed with the pairing results. The other primer pairs were not reliable to the tested isolates due to incongruences in different PCR runs and were not used for mating type determination. The results observed in this study provide additional evidence that the primer pair W16-1/W16-2 (Judelson et al. 1995) is useful and reliable for mating type characterization of the population of *P*. *infestans* in Brazil.

The PCR-based method allows fast mating type determination and does not require pathogen isolation, a very laborious procedure, mainly for tomato isolates. On the other hand, the markers available cannot identify A1/A2 or SF isolates. The A1/A2 and SF isolates are identified as the A1 mating type in this study and also in the study conducted by Zanotta (2019). A new molecular marker able to identify A1/A2 and SF isolates is necessary to replace the pairing completely and assess mating type changes in the population faster.

Isolates able to produce oospore with both mating types or by themselves in Brazil have been reported since 2013 (Casa-Coila et al. 2017; Santana et al. 2013; Zanotta 2019). The occurrence of isolates producing oospore with both mating types was reported in the states of RS (Casa-Coila et al. 2017; Santana et al. 2013) and in SC, in this study. The SF isolates were reported in PR, SP, and MG states (Casa-Coila et al. 2017; Zanotta 2019), The occurrence of A1/A2 and SF individuals increases the chance of recombination by sexual reproduction, as well the co-occurrence of both mating types in fields of MG and PR states reported in the present study and in studies conducted previously (the states of RS, MG, SP, and PR) (Gomes et al. 2007; Oliveira 2010; Zanotta 2019).

The occurrence of sexual reproduction can affect the epidemiology of late blight (Fry and Mizubuti 1998; Mayton et al. 2000; Shattock et al. 1986) because it increases genetic diversity and allows the pathogen to survive for a long time without a host. Thus, given the reports of possible events of sexual reproduction periodical monitoring of the population should be adopted to support the choice of more appropriate late blight management strategies for disease control.

Another sign of alteration in the population is the presence of two genotypes identified in association with potato and a new lineage occurring on tomato. The lineages already reported in Brazil were US-1 on tomato and potato, BR-1 (Brommonschenkel 1988; Goodwin et al. 1994; Reis et al. 2003, 2006; Santana et al. 2013) and EU_2_A1 on potato (Zanotta 2019). Of the genotypes identified in this study, only one has been previously described in Brazil, the EU_2_A1. This was the most commonly found lineage in Europe before 2006, when it was replaced by EU_13_A1, which spread to eastern Africa by imported seeds (Beninal et al. 2022; Puidet et al. 2023). The genotype EU_37_A2, also identified in isolates from potato, was described for the first time in 2013 in the Netherlands and quickly spread in Europe from 2015 (Puidet et al. 2023; Schepers et al. 2018). This genotype increased its frequency and competed well with other lineages (e.g. EU_13_A1) due to characteristics such as resistance to fluazinam, shorter latent period, and higher lesion growth rate (Puidet et al. 2023). The SA_1_A2 genotype, identified in all isolates from tomato and one from potato, has a DNA fingerprint not described before. Although one isolate from potato (Pi01) has been identified as SA_1_A2, the municipality of origin of Pi01 was Viçosa - MG, near an important tomato-producing region and the migration from the surrounding tomato fields to potato plants has been postulated before (Oliveira 2010; Reis et al. 2003). For these reasons, the new genotype seems to be almost exclusively associated with tomato host.

The mtDNA haplotypes Ia and Ib were reported before associated with isolates obtained from potato and tomato (Oliveira 2010). The Ib mtDNA haplotype in isolates of A2 mating type were associated to be resulted from recombination by sexual reproduction or migration of new genotypes of *P*. *infestans* (Oliveira 2010). Isolates of A2 mating type and Ib haplotype were identified in tomato fields associated with the new genotype.

The presence of new genotypes of *P*. *infestans* in Brazil can be explained by the import of infected potato seeds or rare events of recombination. The potato producers in Brazil import seeds tubers mainly from The Netherlands, France, Germany, Canada, and United States (Oliveira 2010). In Europe, the recombination by sexual reproduction was reported in The Netherlands and Nordic countries (Brurberg et al. 2011; Drenth et al. 1994). Also, in Canada the occurrence of sexual recombination was reported recently (Babarinde 2024). Considering the evidences of clonality in the population of *P*. *infestans* in Brazil, the origin of the new SSR genotypes is likely to be due to migrations from European or North American countries, where recombination can take place generating new genotypes. Currently, USA and Canada are major sources of potato seed tubers to Brazil.

Historically, the Brazilian population of *P*. *infestans* is structured by host and previous studies provided evidence for host specialization (Reis et al. 2003, 2006; Santana et al. 2013; Suassuna et al. 2004). In addition, it has been reported that one subpopulation that used to be associated with tomato responds differently from the subpopulation associated with potato regarding the effects of temperature on growth, sporulation, and virulence (Maziero et al. 2009; Suassuna et al. 2004). More recently, phenotypic differences in isolates from tomato were observed: recalcitrance in isolation, low mycelial growth rate on culture medium, long latent period, and long time to zoospore release (data not shown). Considering phenotypic variation can help to forecast pathogen evolution and, consequently, epidemic changes (Ayala-Usma et al. 2020), we hypothesized a shift in the population of *P*. *infestans* in Brazil is occurring. Our hypothesis can be supported by the results presented in this study: (i) the occurrence of A2 mating type in tomato fields for the first time; (ii) a new genotype, the SA_1_A2, associated with isolates from tomato; (iii) the presence of EU_37_A2 genotype in potato fields for the first time, (iv) the occurrence of A1 and A2 mating types in similar ratios in potato fields; and (v) the support of phylogenetic analysis in distinguish the Brazilian isolates from tomato and potato between them.

In the multigene phylogenetic analysis, the distinction between isolates of the SA_1_A2 genotype (tomato host) and the other isolates (potato host) is well supported. This result can add to the body of evidence of host specialization of *P*. *infestans* in Brazil. In the phylogenetic analysis of the single genes, the tree for G3PDH provides the most convincing evidence for isolate divergence according to the host of origin. This gene has an important function in the initial stages of plant-pathogen interactions (Laxalt et al. 1996; Shan et al. 2004). The phylogenetic difference between isolates from both hosts can explain the phenotypic variation observed in isolates obtained from tomato such as longest latent period, and longest time to zoospore release compared to isolates from potato.

The evidence of change in genetic profile of the population of *P*. *infestans* in Brazil can be well visualized by the temporal analysis of the population from 1998 to 2024. The analysis of the 1998-2010 and 2020-2024 subpopulations show the contrast of genetic composition of the Brazilian population over the years. The higher diversity of the 1998-2010 subpopulation can be explained by migration of new genotypes and by the sexual recombination events evidenced by the index of association and detected occurred between 2003 and 2005 (Oliveira 2010).

The population of *P*. *infestans* from Brazil, despite predominant asexual characteristics, has undergone changes over the years. This study characterized the current Brazilian population of *P*. *infestans* mating type, mitochondrial DNA haplotypes, 12-plex SSR genotype, and mefenoxam sensitivity. For the first time, the A2 mating type is associated with isolates from tomato in Brazil. Also, three genotypes are co-occurring in the population and a new lineage seems to occur exclusively in tomato fields. Phylogenetic evidence of two host-defined populations supported the phenotype differences observed in isolates from tomato and potato. The update of the Brazilian population of *P*. *infestans* and the separation of isolates from potato and tomato are very important information about this pathogen in Brazil that should be considered to take adequate late blight management actions to each host.

## Acknowledgements

The authors express their gratitude to Fundação de Amparo à Pesquisa do Estado de Minas Gerais - FAPEMIG, by grant number APQ-01387-08, and a Coordenação de Aperfeiçoamento de Pessoal de Nível Superior – Brasil (CAPES) – Financing code 001” for partially funding the current research.

## Supplementary material

**Table S1.** SSR allele names and sizes adapted from Li et al. (2013) to Brazilian isolates of *Phytophthora infestans* according control isolates sent by Dr. David Cooke (JHI).

|  |  |  |  |  |  |  |  |  |  |  |  |  |  |  |  |  |  |  |  |
| --- | --- | --- | --- | --- | --- | --- | --- | --- | --- | --- | --- | --- | --- | --- | --- | --- | --- | --- | --- |
| G11 | Allele name | 130 | 132 | 138 | 140 | 142 | 146 | 148 | 150 | 152 | 154 | 156 | 158 | 160 | 162 | 164 | 166 | 168 | 170 |
|  | Allele size | 130.3 | 132.3 | 138.5 | 140 | 142 | 147.1 | 149.4 | 151.4 | 153.1 | 155.2 | 157 | 159.4 | 161.1 | 163.1 | 165.1 | 167 | 169.2 | 171.4 |
|  | Allele name | 172 | 176 | 198 | 200 | 202 | 204 | 206 | 208 | 210 | 214 | 218 | 220 | 222 |  |  |  |  |  |
|  | Allele size | 173.2 | 176.7 | 198.3 | 200.2 | 202 | 204.1 | 206 | 207.7 | 210 | 213.6 | 217.5 | 219.5 | 221.4 |  |  |  |  |  |
| Pi04 | Allele name | 160 | 166 | 168 | 170 | 172 |  |  |  |  |  |  |  |  |  |  |  |  |  |
|  | Allele size | 167 | 171.2 | 173.5 | 175.4 | 179 |  |  |  |  |  |  |  |  |  |  |  |  |  |
| Pi4B | Allele name | 203 | 205 | 209 | 211 | 213 | 215 | 217 | 221 | 225 |  |  |  |  |  |  |  |  |  |
|  | Allele size | 206 | 207 | 210.5 | 212.5 | 215 | 218 | 220 | 224 | 229 |  |  |  |  |  |  |  |  |  |
| Pi63 | Allele name | 270 | 273 | 279 |  |  |  |  |  |  |  |  |  |  |  |  |  |  |  |
|  | Allele size | 272 | 275 | 280 |  |  |  |  |  |  |  |  |  |  |  |  |  |  |  |
| Pi70 | Allele name | 189 | 192 | 195 | 198 |  |  |  |  |  |  |  |  |  |  |  |  |  |  |
|  | Allele size | 187 | 191 | 194 | 196 |  |  |  |  |  |  |  |  |  |  |  |  |  |  |
| D13 | Allele name | 108 | 110 | 112 | 114 | 116 | 118 | 120 | 122 | 124 | 126 | 128 | 130 | 132 | 134 | 136 | 138 | 140 | 142 |
|  | Allele size | 106.2 | 108.2 | 110 | 112.1 | 114.2 | 116.3 | 119 | 120.9 | 123.1 | 125.1 | 127.1 | 128.8 | 130.6 | 132.6 | 135 | 137 | 138.9 | 141.1 |
|  | Allele name | 144 | 146 | 148 | 150 | 152 | 154 | 156 | 158 | 160 | 162 | 164 | 166 | 168 | 170 | 172 | 174 | 176 | 178 |
|  | Allele size | 143.3 | 145.6 | 147.8 | 150.2 | 152.3 | 154.3 | 156.4 | 158.6 | 160.6 | 162.4 | 164.4 | 166.6 | 168.4 | 170.4 | 172.5 | 174.4 | 176.6 | 178 |
|  | Allele name | 184 | 188 | 190 | 208 | 210 | 212 | 214 | 216 |  |  |  |  |  |  |  |  |  |  |
|  | Allele size | 185.1 | 188.9 | 190.8 | 208.7 | 210.8 | 212.7 | 214.7 | 217 |  |  |  |  |  |  |  |  |  |  |
| SSR2 | Allele name | 163 | 167 | 173 | 175 | 177 |  |  |  |  |  |  |  |  |  |  |  |  |  |
|  | Allele size | 163.5 | 167.3 | 173.3 | 175.4 | 177.4 |  |  |  |  |  |  |  |  |  |  |  |  |  |
| Pi02/SSR3 | Original allele name | 152 | 158 | 160 | 162 | 164 | 166 | 168 |  |  |  |  |  |  |  |  |  |  |  |
|  | Allele name | 258 | 264 | 266 | 268 | 270 | 272 | 274 | 262 |  |  |  |  |  |  |  |  |  |  |
|  | Allele size | 260 | 266 | 268 | 270 | 272 | 274 | 276 | 264.7 |  |  |  |  |  |  |  |  |  |  |
| SSR4 | Allele name | 283 | 285 | 287 | 289 | 291 | 293 | 295 | 297 | 299 | 301 | 303 | 305 | 307 | 309 | 311 | 313 | 315 | 317 |
|  | Allele size | 281.4 | 285 | 288 | 289 | 290 | 292 | 294 | 296 | 298 | 299.7 | 301.9 | 304 | 305.7 | 307.9 | 309.9 | 312.1 | 314 | 316.3 |
| SSR6 | Allele name | 232 | 236 | 240 | 242 | 244 | 246 | 254 | 256 | 258 | 260 | 262 |  |  |  |  |  |  |  |
|  | Allele size | 234 | 236.1 | 245 | 245 | 247 | 249 | 251 | 253 | 255 | 259.8 | 262.7 |  |  |  |  |  |  |  |
| SSR8 | Allele name | 260 | 264 | 266 |  |  |  |  |  |  |  |  |  |  |  |  |  |  |  |
|  | Allele size | 262 | 266 | 268 |  |  |  |  |  |  |  |  |  |  |  |  |  |  |  |
| SSR11 | Allele name | 331 | 341 | 356 |  |  |  |  |  |  |  |  |  |  |  |  |  |  |  |
|  | Allele size | 331 | 341 | 356 |  |  |  |  |  |  |  |  |  |  |  |  |  |  |  |

**Supplementary 2.**
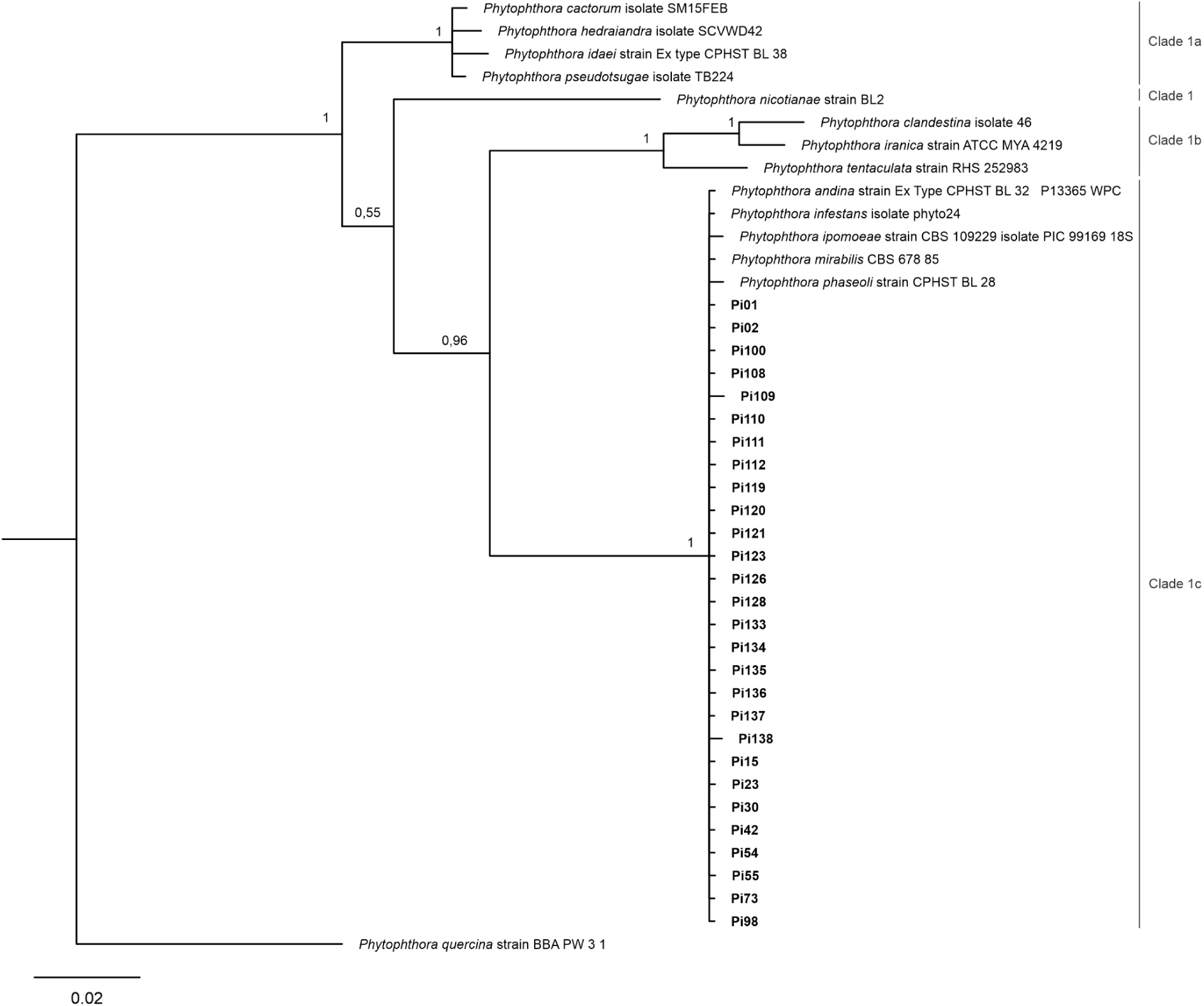
Bayesian 50 % majority rule tree based on the *ITS* loci of 27 Brazilian isolates of *Phytophthora infestans* obtained from potato (n = 11) and tomato (n = 16) and reference sequences retrieved from GenBank (isolate description shown in the tree). Bayesian posterior probabilities (%) were also calculated and values are shown at each node > 0.50. The tree was rooted in *P*. *quercina* (Clade 3). The scale bar represents the expected number of substitutions per site. The Brazilian isolates used in the analyses are indicated by their code in bold. The *Phytophthora* clade of isolates are indicated by bar and the respective clade name.

**Supplementary 3.**
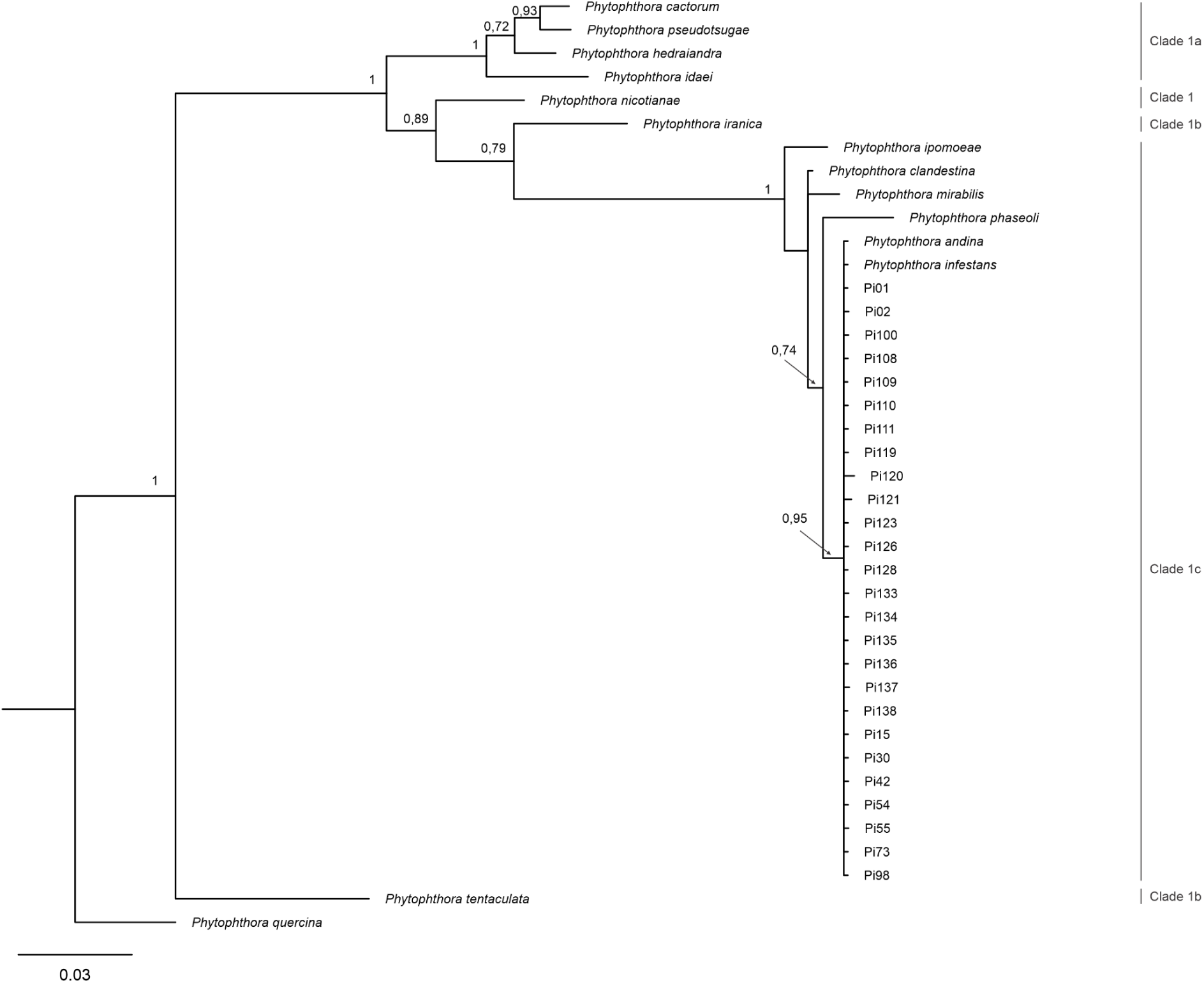
Bayesian 50 % majority rule tree based on the *CoxI* loci of 27 Brazilian isolates of *Phytophthora infestans* obtained from potato (n = 11) and tomato (n = 16) and reference sequences retrieved from GenBank. Bayesian posterior probabilities (%) were also calculated and values are shown at each node > 0.70. The tree was rooted in *P*. *quercina* (Clade 3). The scale bar represents the expected number of substitutions per site. The Brazilian isolates used in the analyses are indicated by their code in bold. The *Phytophthora* clade of isolates are indicated by bar and the respective clade name.

